# Two-dimensional HRS condensates drive the assembly of flat clathrin lattices on endosomes

**DOI:** 10.1101/2024.10.01.616068

**Authors:** Markku Hakala, Satish Babu Moparthi, Iva Ganeva, Mehmet Gül, Javier Espadas, Carlos Marcuello, César Bernat-Silvestre, Adai Colom, Mikhail Kudryashev, Wanda Kukulski, Stéphane Vassilopoulos, Marko Kaksonen, Aurélien Roux

## Abstract

In cells, the curved clathrin structures in vesicle budding are well characterized, while the flat ones remain poorly understood. We reconstituted the flat assembly of ESCRT-0 protein HRS and clathrin onto lipid membranes *in vitro*. HRS was found to form protein condensates. These condensates spread as a two-dimensional layer on negatively charged membranes and promoted the assembly of clathrin into a flat coat. Correlative cryo-tomography of HRS-labeled endosomes revealed a pure hexagonal lattice, consistent with flat clathrin structures. Cholesterol enhanced HRS recruitment to the membrane both in cells and in supported bilayers. Furthermore, cholesterol promoted the phase separation of HRS onto membranes, which in turn concentrated cholesterol underneath. This positive feedback promoted the formation of HRS-clathrin microdomains that sorted reconstituted ubiquitinated cargoes. Altogether, our results show that the unique architecture of ESCRT-0 is assembled by the two-dimensional phase-separation of HRS which drives the assembly of flat clathrin coats.

## Introduction

Clathrin, the main component of the coat involved in clathrin-mediated endocytosis (CME), self-assembles i*n vitro* into spherical cages that resemble the coats observed on endocytic vesicles (Ybe et al., 1999; Liu et al., 1995; Roth and Porter, 1964; Ungewickell and Branton, 1981; Crowther and Pearse, 1981). During CME, membrane-binding adaptor proteins drive clathrin assembly into hexagons and pentagons, promoting membrane deformation and invagination (Kaksonen and Roux, 2018; Sochacki and Taraska, 2019). Besides CME, clathrin is involved in intracellular vesicle trafficking routes between endosomes, trans-Golgi network, and the plasma membrane. During these trafficking processes, the clathrin coat assembles on the organellar membrane and facilitates membrane budding in a similar way to CME (Vassilopoulos and Montagnac, 2024). However, how adaptors control the position and the ratio between pentagons and hexagons, setting membrane curvature, remains unknown.

In addition to vesicular clathrin structures that promote membrane deformation, mammalian cells have flat and tubular clathrin structures, which are usually not linked to vesicle trafficking (Vassilopoulos and Montagnac, 2024). At the plasma membrane, hexagonal clathrin lattices participate in the formation of integrin-dependent adhesion plaques or collagen-pinching tubular adhesions (Lock et al., 2019; Hakanpää et al., 2023; Elkhatib et al., 2017). Among the flat clathrin lattices, the least characterized are found on endosomes. There, clathrin associates with the endosomal sorting complex required for transport (ESCRT)-0 by binding with Hepatocyte growth factor tyrosine kinase substrate HRS/HGS (from now on, HRS) (Raiborg et al., 2001a; Sachse et al., 2002). HRS-clathrin domains sort ubiquitylated cargoes, which are further concentrated in intraluminal vesicles (ILV) (Raiborg et al., 2002; Wenzel et al., 2018; Migliano et al., 2023). Remarkably, HRS-clathrin coats have a distinctive bi-layered organization (Sachse et al., 2002), which is unique among clathrin structures in the cell (Vassilopoulos and Montagnac, 2024). However, how the unique architecture of the ESCRT-0-clathrin coat is assembled on endosomes and how it is linked to its sorting function remains unanswered.

Cholesterol is more abundant in membranes where clathrin-mediated trafficking occurs: the trans-Golgi network, endosomes, and the plasma membrane (Kim and Burd, 2023; Bigay and Antonny, 2012). Cholesterol plays a crucial role in lipid phase separation and sets membrane biophysical properties, making it essential for numerous membrane functions. For example, cholesterol promotes membrane bending during CME (Anderson et al., 2021; Rodal et al., 1999; Pichler and Riezman, 2004). Cholesterol is highly enriched in multivesicular endosomes (Van Meer et al., 2008; Ikonen, 2018), and its trafficking in cells is tightly regulated (Wheeler and Sillence, 2020; Ikonen and Zhou, 2021). An imbalance of this tight regulation causes diseases, as the excess of cholesterol in endosomes alters transport functions in the Niemann-Pick type C disease caused by mutations in *NPC1* or *NPC2* genes (Carstea et al., 1997; Naureckiene et al., 2000). While its endosomal excess causes trafficking defects, the physiological roles of cholesterol at endosomes remain poorly characterized. Furthermore, while both cholesterol and clathrin are enriched in the same membrane compartments, whether the cholesterol and clathrin assembly impact each other has not been investigated.

Here, we studied the assembly of ESCRT-0-clathrin coats on endosomes and their interplay with the underlying membrane. Our reconstitution approach showed that HRS recruitment to the membrane depended on its interaction with PI(3)P and was enhanced by cholesterol. HRS, which contains unstructured domains, formed spherical condensates in solution and two-dimensional condensates on membranes. Two-dimensional HRS condensates promoted the assembly of a solid and dense, flat coat of clathrin. In cells, the recruitment of HRS and clathrin on endosomes also depended on cholesterol, and cryo-electron tomography showed that clathrin forms a flat, hexagonal protein lattice.

## Results

### HRS recruits clathrin on membranes as flat lattices and domes

Since clathrin does not directly bind to membranes, the formation of various clathrin structures on cellular membranes relies on organelle-specific adaptors, with HRS being uniquely associated with endosomes. Its FYVE domain binds phosphatidyl-inositol 3-phosphate (PI(3)P), which is enriched in early endosomal membranes and intraluminal vesicles (Gillooly, 2000; Raiborg et al., 2001b; Urbé et al., 2000). To understand their assembly mechanism, we initially aimed at reconstituting the ESCRT-0 clathrin coat *in vitro* using purified and labeled human HRS and bovine clathrin. We measured the dynamics of clathrin recruitment on HRS-coated supported lipid bilayer (SLB) membranes containing 1mol% PI(3)P, 44mol% DOPC, 20mol% DOPS, 20mol% DOPE, and 15mol% cholesterol (Lipid mix 1, Table S2) by using microfluidics-assisted sequential addition of proteins (Figure 1A-B). Our experiments revealed that HRS recruits clathrin on SLBs in a concentration-dependent manner (Figure 1C-E) with both the nucleation rate and the final amount of membrane-bound clathrin proportional to the bulk HRS concentration (Figure 1D-E). Clathrin recruitment followed a single exponential kinetics but did not saturate at the high HRS concentrations.

**Figure 1.**
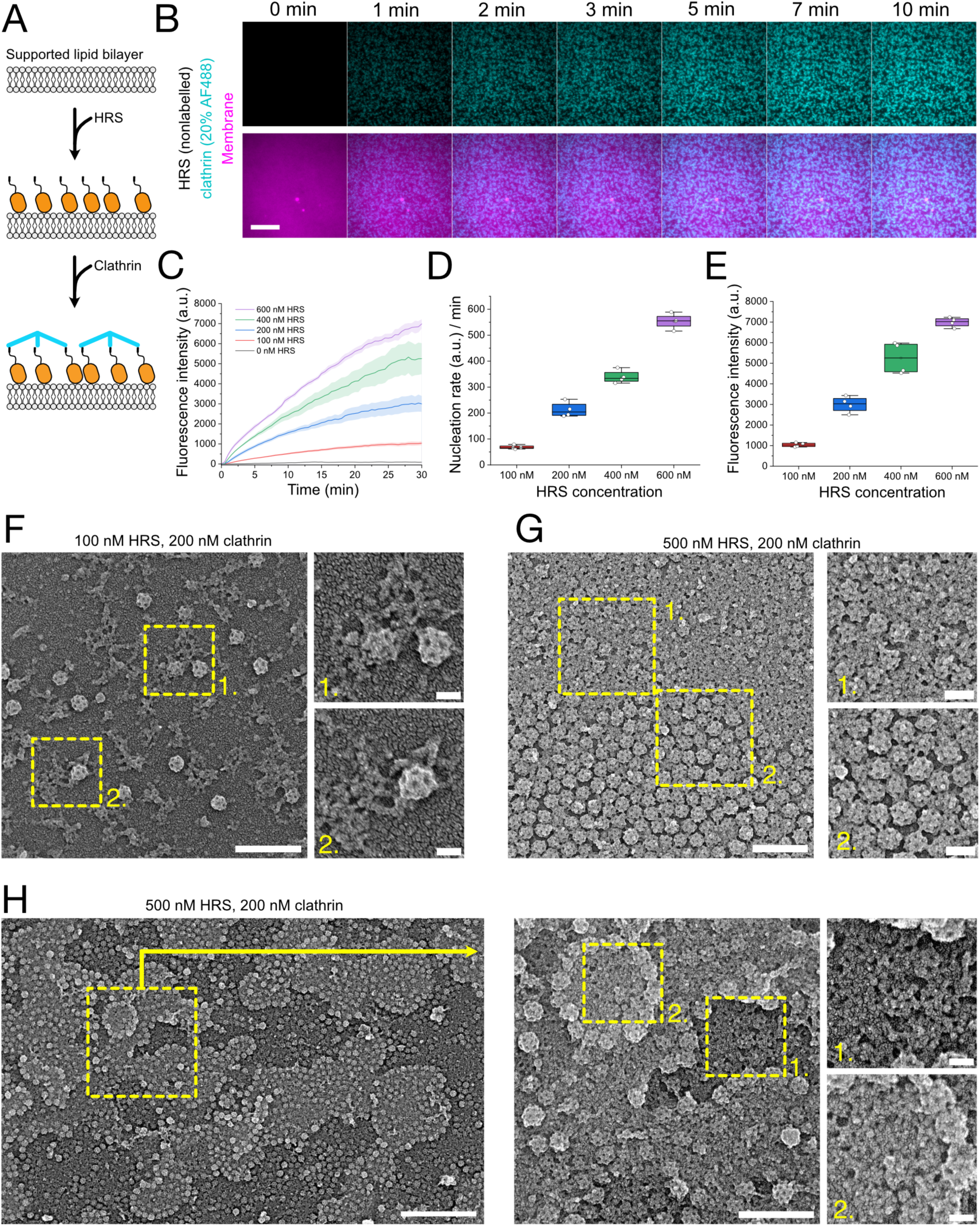
HRS recruits clathrin on membranes as flat patches and spherical cages. (A) A schematic illustration of the reconstitution strategy on SLBs using sequential protein addition with a microfluidics device. (B) The recruitment of 200 nM clathrin (20% AlexaFluor-488 labeled, cyan) over time on SLBs (magenta) preincubated with 500 nM nonlabelled HRS. The scale bar is 10 µm. (C) The recruitment of 200 nM clathrin (20% AlexaFluor-488 labeled) over time with different HRS concentrations. Data is the mean of four experiments with standard deviations shown. (D-E) Nucleation rates (D) and final clathrin fluorescence intensities (E) plotted over HRS concentration from the data in panel (B). (F-H) Representative PREM images of reconstituted HRS-clathrin lattices on SLBs. Two different HRS concentrations were used: (F) 100 nM HRS and (G-H) 500 nM HRS with clathrin concentration being 200 nM in both samples. Scale bars in panels (F-G) for larger images are 300 nm and for zoom-in images 50 nm. Scale bars in panel (H) are 1 µm for the left image, 300 nm for the first zoom-in, and 50 nm for the second zoom-in image.

Next, we aimed to better understand the recruitment of HRS on membranes. As expected, HRS and clathrin were recruited onto PI(3)P-rich giant unilamellar vesicle (GUV) membranes (Lipid mix 1, Table S2), but not in the absence of PI(3)P (Lipid mix 2, Table S2) (Figure S1A-B). To test the recruitment kinetics of HRS onto PI(3)P-rich membranes, we prepared SLBs with 1% PI(3)P (Lipid mix 1, Table S2), and followed the recruitment of labeled HRS by total internal reflection fluorescence (TIRF) microscope. Recruitment of HRS followed single exponential kinetics, saturating after 15-20 minutes with 100 and 250 nM bulk HRS and significantly faster with 500 nM bulk HRS (Figure S1C). 100 nM bulk HRS led to a protein coat on the membrane that showed moderate fluorescence recovery after photobleaching, an indication of lateral protein movement on the membrane after recruitment. In contrast, at 250 nM and 500 nM bulk HRS, the recruitment led to a stable protein coat with minimal lateral movement (Figure S1D).

Our reconstitution experiments indicated that HRS promotes clathrin recruitment in a concentration-dependent manner, and we wanted to visualize these HRS-clathrin coats at a higher resolution. Platinum replica electron microscopy (PREM) is a powerful approach to visualize protein lattices on flat surfaces because it provides high contrast and shadowing, suited to visualize flat polygonal lattices (Heuser, 1980; Wernert et al., 2024; Sochacki et al., 2017). We deposited SLBs on a glass substrate and compared clathrin recruitment kinetics to the resulting lattice morphology. We prepared samples by incubating SLBs first for 60 minutes with 100 nM or 500 nM concentrations of HRS, then washed five times with reaction buffer to remove the remaining bulk HRS and finally incubated with 200 nM clathrin for 30 minutes. Samples were then fixed and platinum replicas were prepared for electron microscopy analysis. With 100 nM HRS, we observed small, flat clathrin patches on SLBs (Figures 1F and S2A). These patches were often accompanied by spherical clathrin cages. Because these cages were always connected to a neighboring clathrin patch (Figure 1F), we reasoned that they originated from initially flat clathrin lattices or were assembled as cages next to flat protein patches. With 500 nM HRS and 200 nM clathrin, we frequently observed large patches of flat and dense clathrin islands along with regions where clathrin formed cup-shaped structures (Figure 1G). Notably, these cup-shaped structures appeared to have various degrees of curvature (Figure 1G).

We often observed round clathrin islands in our PREM samples prepared with 500 nM of HRS and 200 nM of clathrin (Figures 1H and S2B). These clathrin islands had a dense and predominantly flat appearance. They appeared as stacked flat patches coated with clathrin, having multiple layers of similar thickness (Figure S2C-D). However, clathrin looked flatter on the layer closer to the glass. The observation of flat clathrin patches raised the question of how they are formed. Their appearance suggests they form on top of protein or membrane domains, which are surrounded by more curved clathrin coat regions: Indeed, tilt series of these samples supported that the clathrin patches are thicker than just the clathrin layer (Movie S1).

We reasoned that a specific mechanism was needed to form such defined and round-shaped clathrin patches. Intrigued by the disordered nature of HRS (AlphaFold Protein Structure Database, AF-O14964-F1-v4), we hypothesized that HRS might phase-separate to form protein patches onto which clathrin could assemble. Supporting this notion, we occasionally observed highly dense protein domains on GUVs and SLBs when these membranes were incubated with 500 nM of HRS (Figure S1E-F). Notably, the size of these protein domains was in the same range as the clathrin patches observed by PREM (Figures 1H and S2B). To understand the nature of the HRS domains onto which flat clathrin lattices were formed, we further investigated if HRS could undergo phase separation in solution and on membranes.

### HRS forms condensates that wet the supported lipid bilayer membranes

According to UIPred3 predictions (https://iupred3.elte.hu), HRS contains two Intrinsically Disordered Regions (IDRs): one between the DUIM motif and the helical region and the other one at its carboxy terminus (Figure S3A). Intrinsically disordered regions (IDRs) often drive liquid phase separation of proteins into condensates (Brangwynne et al., 2015). Consistently with the high proportion of disordered regions in its sequence, we observed that labeled HRS at 2 µM formed liquid droplets of protein condensates (Figure S3B). These droplets could grow by fusion and coalescence, supporting their liquid nature (Movie S2), a defining characteristic of phase-separated protein condensates (Banani et al., 2017).

We wondered how these condensates would interplay with membranes. To test this, we introduced 2 µM labeled HRS on SLBs containing 1% PI(3)P (Lipid mix 1, Table S2). The protein condensates interacted with the membrane and started spreading on it (Figure 2A, orange arrows). Protein droplets spread entirely on the membrane surface and fused into a continuous and homogenous layer of HRS fully covering the membrane (Figure 2B, orange arrows) (Movie S3), further indicating that HRS condensates are liquid. This process resembled surface wetting by liquid droplets. The fluorescence intensity of HRS remained rather constant throughout the wetted area, indicating that the HRS layer that spreads onto the membrane forms a two-dimensional film (Figure 2C). We also noticed the spontaneous formation of HRS domains with the same intensity than the two-dimensional condensate growing out from droplets (Figure 2A-B, yellow arrowheads). This suggested that the two-dimensional condensate can form from the low-density phase of HRS on the membrane surface. To test if protein can diffuse in HRS condensates on the membrane, we photobleached small regions of the droplet-like condensates, and of the condensate film spread on the membrane (Figure S3C-D). Similarly to the HRS coat assembled with 250-500 nM protein (Figure S1D), neither of these HRS condensate populations showed a fluorescence recovery in the photobleached region within the 3-minute acquisition period (Figure S3C-D). This suggests that when these condensates are spreading on the membrane in a time scale of 1-3 hours (Figure 2A-B), they become very viscous, which would lead to negligible lateral protein diffusion.

**Figure 2.**
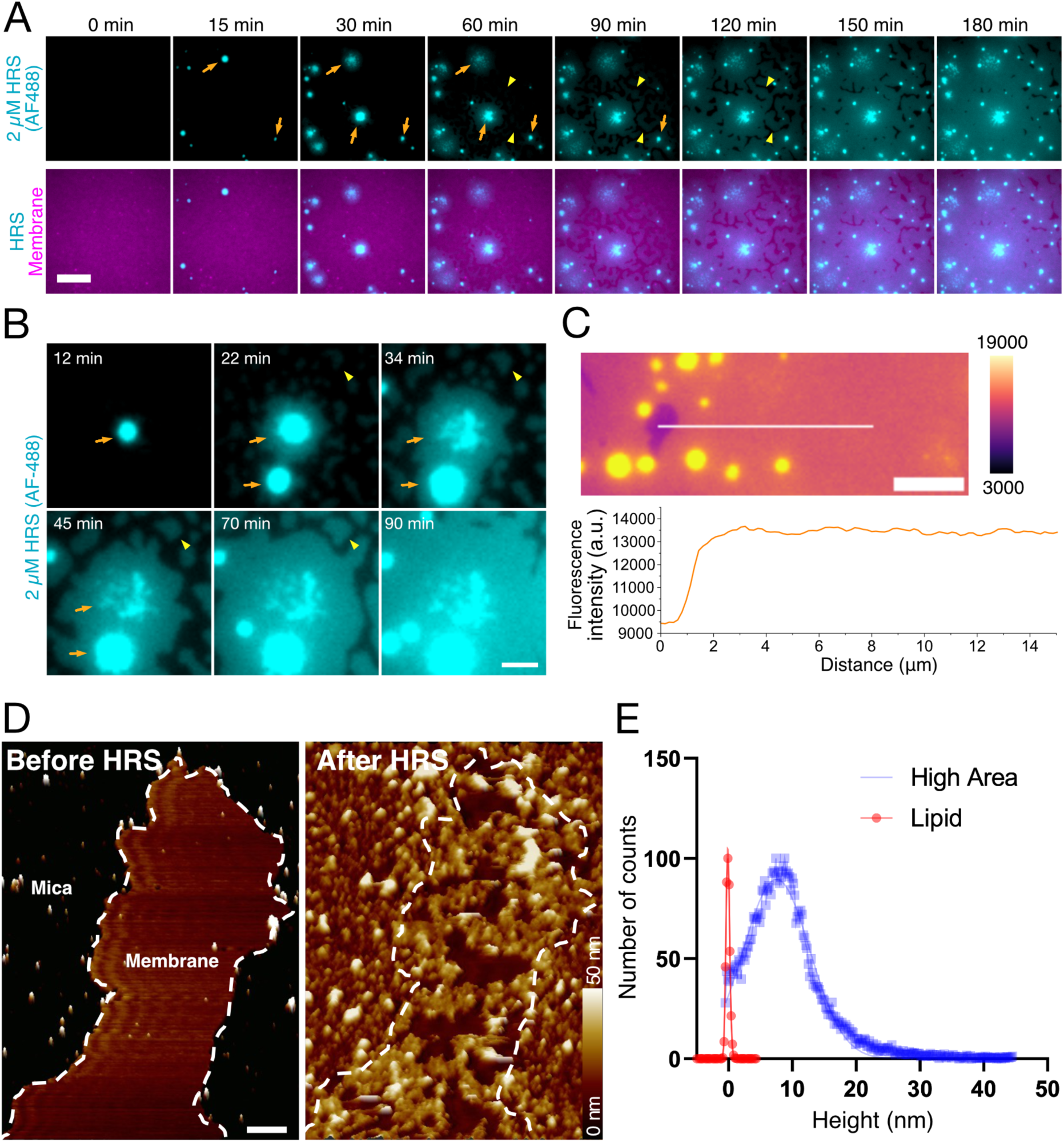
HRS forms two-dimensional condensates on membranes. (A) Representative fluorescence timelapse images of 2 µM AlexaFluor-488 labeled HRS (cyan) on SLBs (magenta). The scale bar is 10 µm. (B) A representative example of HRS condensate droplets (cyan) spreading on the membrane (not shown) over time and droplets getting dispersed. The scale bar is 5 µm. In panels (A-B), orange arrows indicate regions where droplet-like condensates spread on SLBs to form a two-dimensional film, and yellow arrowheads indicate regions where HRS spontaneously forms a two-dimensional film without initial droplet condensate. (C) A fluorescent microscopy image of HRS condensate wetting the membrane with a fluorescence intensity heatmap and a line profile analysis of HRS fluorescence intensity over the condensate. A scale bar is 10 µm. (D). Representative HS-AFM images of HRS condensates on SLBs. Scan rate and resolution were settled at 15 Hz and 512 pixels per frame, respectively. Scale bar is 500 nm. (E) A histogram of HRS condensate thickness on SLB. Each event represents a pixel on SLB.

To better understand the formation of HRS condensates on membranes, we used high-speed atomic force microscopy (HS-AFM). When HRS was injected into the imaging chamber containing SLBs on mica, we initially observed a homogeneous binding of HRS, which separates rapidly into patches (Figure 2D). Notably, these protein patches did not show high-ordered structure, which indicates they form through phase separation (Figure 2D). HRS patches grew over time with larger protein patches being less mobile (Movie S4), supporting the low protein diffusion observations done by FRAP (Figures S1D and S3C-D). HS-AFM also allowed us to measure the thickness of the HRS patches with the height around 10 nm (Figure 2E).

High-energy interactions with membranes are known to favor complete membrane wetting by protein condensates (Zhao et al., 2021). Thus, the strong interaction between HRS and lipids in the SLBs could promote wetting behavior. We tested this by observing whether the protein condensates spread on a membrane composed of DOPC lipids only (uncharged membrane, Lipid mix 7, Table S2) or a membrane containing negatively charged DOPS as well as PI(3)P (Lipid mix 1, Table S2). HRS condensates did not adhere and spread on the DOPC-only membranes but readily spread on the DOPS/PI(3)P-rich membrane as well as membranes containing 20% DOPS but no PI(3)P (Lipid mix 2, Table S2), forming a dense protein film (Figure S3E). Therefore, we concluded that electrostatic interactions of HRS with charged lipids were essential for membrane wetting and growth of the two-dimensional condensate.

HRS droplets fully wetting the membrane resembled the two-dimensional condensates reported in previous studies with other proteins (Lee et al., 2019; Huang et al., 2016; Banjade and Rosen, 2014). However, the physiological functions of such two-dimensional condensates remained speculative. In our system, we could directly test the role of two-dimensional condensates on clathrin assembly.

### Clathrin assembly varies depending on the type of HRS condensates it binds to

We wondered how HRS condensates modify the structure and dynamics of clathrin assembly compared to the low density of HRS on the membrane. To this end, we incubated PI(3)P-rich SLBs (Lipid mix 1, Table S2) with 2 µM HRS. Immediately after droplet-like condensates formed, we washed the remaining HRS out five times with reaction buffer and introduced 200 nM of clathrin. Droplet-like HRS condensates readily recruited clathrin, which did not enter inside the condensates but covered the surface of the protein condensate (Figure S4A). To investigate how clathrin binds to two-dimensional condensates, we used the same protocol but waited until the spreading of two-dimensional condensates occurred before injecting clathrin. Fluorescent clathrin was recruited both onto HRS droplets that had not fully spread into two-dimensional condensates, as well as onto two-dimensional condensates (Figure S4B). In confocal fluorescence microscopy images, we observed that HRS fluorescence intensity increased stepwise from the low-density region to high-density two-dimensional condensate and dramatically increased at the droplet-like condensate (Figure S4B-C). Clathrin fluorescence intensity followed the same trend, increasing at the border of two-dimensional HRS condensates and substantially increasing at the HRS droplet (Figure S4B-C).

Our fluorescence microscopy data with HRS condensates and labeled clathrin support the idea of three different HRS populations on the membrane: 1) low-density HRS, 2) two-dimensional HRS condensates, and 3) droplet-like HRS condensates, which all readily recruit clathrin (Figure 3A). We looked at the clathrin coat architecture on droplets and two-dimensional condensates using PREM. Platinum replica samples prepared with 2 µM HRS and 200 nM clathrin at room temperature showed extensive amounts of clathrin cages at the surface of HRS droplets (Figures 3B and S4D-F). Clathrin structures at the surface of two-dimensional condensates next to the droplet condensates appeared flatter and less numerous than on droplets (Figures 3B-C and S4F-H). Furthermore, we observed areas where flat clathrin lattices were visible (Figures 3C and S4G-H). These flat clathrin lattices were similar to those we observed at 500nM HRS concentration (Figure 1G), supporting the hypothesis that these dense, round-shaped patches of flat clathrin are assembled on two-dimensional HRS condensates.

**Figure 3.**
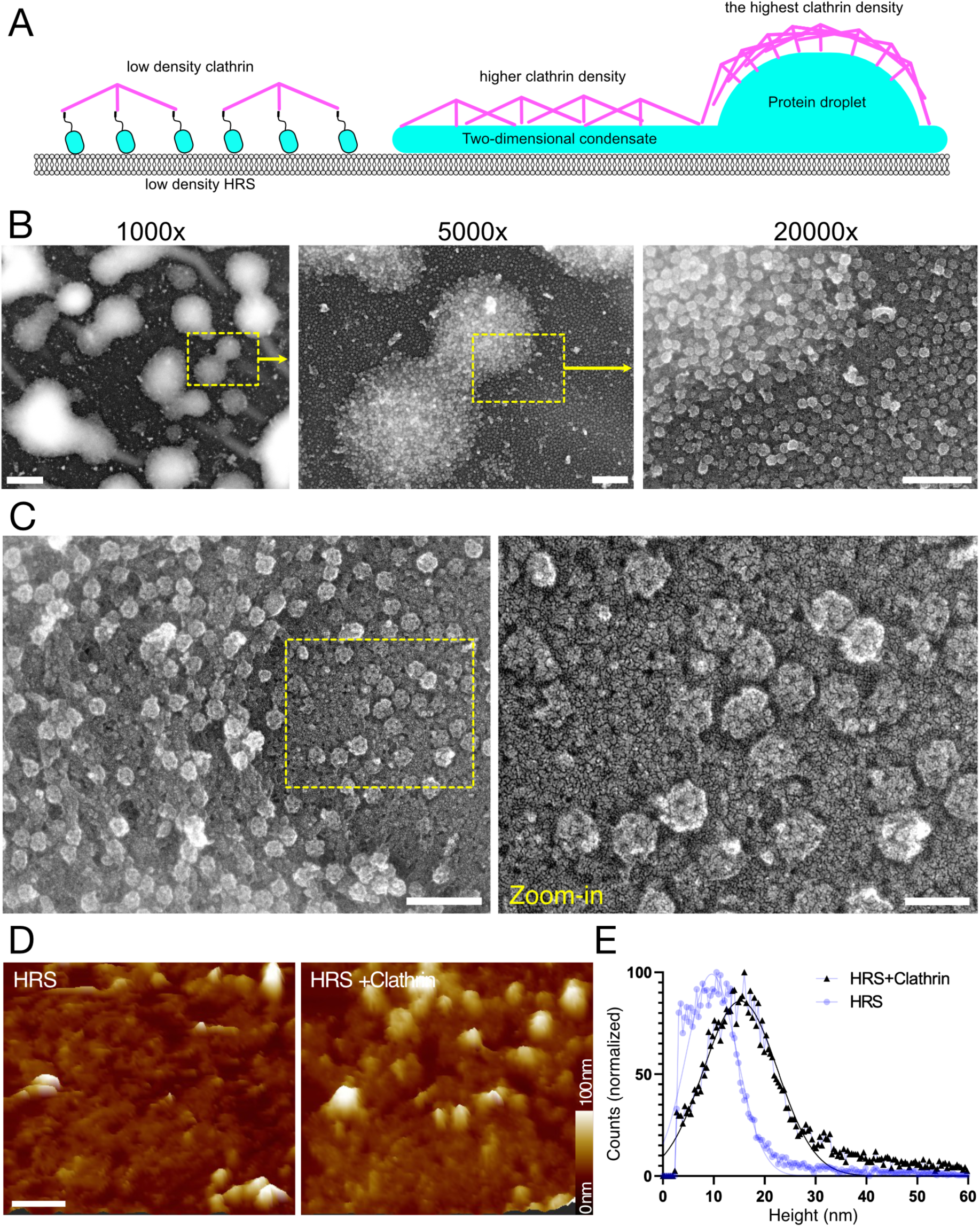
Clathrin assembles on HRS condensates. (A) A schematic illustration of three HRS populations: 1) low-density patches, 2) a two-dimensional condensate, 3) droplet condensate, and how clathrin assembles onto these HRS populations. (B) Representative PREM images of reconstitution samples with 2 µM HRS and 200 nM clathrin. Scale bars from left to right: 5 µm, 1 µm, 500 nm. (C) Representative platinum replica microscopy images of 2 µM HRS and 200 nM clathrin. Scale bars are 300 nm for the larger image and 100 nm for the zoom-in. (D) Representative HS-AFM images of HRS and HRS-clathrin coats on SLBs. The scale bar is 500 nm. (E) A histogram of HRS and HRS-clathrin coat heights. Each event represents a pixel on SLB.

We then wanted to measure the height of HRS-clathrin coat. To this end, we reconstituted 500 nM HRS and clathrin on SLBs on the Mica surface and imaged them with HS-AFM (Figure 3D, Movie S5). As earlier, the thickness of HRS condensates was around 10 nm (Figure 3E). When clathrin was assembled on these two-dimensional HRS condensates, it yielded a coat with a median height of 16-18 nm (Figure 3E). Consistent with our observations of different layers of HRS-clathrin (Figures 3A and S4B-C), HRS-clathrin coats imaged with HS-AFM had regions with different heights (Figure S5A-B). The height of the thicker regions was around 20-30 nm, while the height of the thinner ones was 15-20 nm, consistent with stacking of layers (Figure S5A-B). We observed hexagonal and pentagonal clathrin structures especially on the thicker regions, while the thinner regions appeared to contain less structured clathrin coat (Figure S5C-D). This supports the notion that the HRS-clathrin assembly is multi-layered, and that the clathrin is assembled into flat coats primarily on the thinner regions, corresponding to a single, two-dimensional HRS condensate.

Taken together, our experiments revealed that clathrin readily assembles onto HRS condensates. Two-dimensional condensates promote the assembly of both flat, dense clathrin coats as well as spherical cages (Figures 1H, 2C-D, and S2B). On the other hand, HRS droplets promoted clathrin assembly into hexagonal and pentagonal spherical cages (Figure 3B). We next wondered how similar these *in vitro* clathrin assemblies mediated by condensation of HRS were with the in vivo ESCRT-0 assemblies found on endosomes.

### Correlative cryo-electron tomography reveals the three-layered protein coat and a hexagonal clathrin lattice on endosomes

Previous studies have identified flat clathrin microdomains on endosomes by electron microscopy of resin-embedded cells (Raiborg et al., 2002; Sachse et al., 2002). However, these studies did not visualize the clathrin coat architecture. We aimed to reveal endosomal clathrin lattices using cryo-electron tomography (cryo-ET) combined with subtomogram averaging, a set of methods to image and structurally analyze molecules in situ. To localize HRS-clathrin structures in cells, we tagged endogenous HRS in HeLa MZ cells with mScarlet-I3 fluorescent protein (see Methods). We confirmed the correct localization of mScarlet-I3-HRS by labeling endosomes and lysosomes with a pulse of AlexaFluor-647 labeled epidermal growth factor (AF647-EGF), which is commonly used to detect different compartments of intracellular vesicle trafficking (Wenzel et al., 2018). Endogenously tagged mScarlet-I3-HRS colocalized with AF647-EGF after 15 minutes of AF647-EGF incubation, indicating that mScarlet-I3-tagged HRS localizes to endosomes (Figure S6C-D). These results confirmed that mScarlet-I3 tagging did not affect the localization of HRS in our cell line.

We then used cryo-correlative light and electron microscopy (cryo-CLEM) to acquire cryo-ET data of HRS-positive endosomes (Figure S7A-D) in our Scarlet-I3-HRS HeLa cells and in a previously described HeLa cell line stably expressing mCherry-HRS (Wenzel et al., 2018; Migliano et al., 2023). In the resulting tomograms of target areas containing either mCherry-HRS, mScarlet-I3-HRS or Alexa-Fluor-647 labeled EGF signals, we observed patches of a dense protein coat on endosomes (Figures 4A and S7E-P). This coat extended approximately 40 - 60 nm from the membrane (Figure 4A, yellow brackets). Consistent with previous reports (Sachse et al., 2002; Raiborg et al., 2002), we found these protein coats to have a flat and multi-layered appearance, very similar to the structures we observed in our reconstituted samples by PREM and HS-AFM (Figure 4A-C). We observed a particularly dense layer positioned 15-20 nm away from the membrane (Figure 4B-C, yellow arrows).

**Figure 4.**
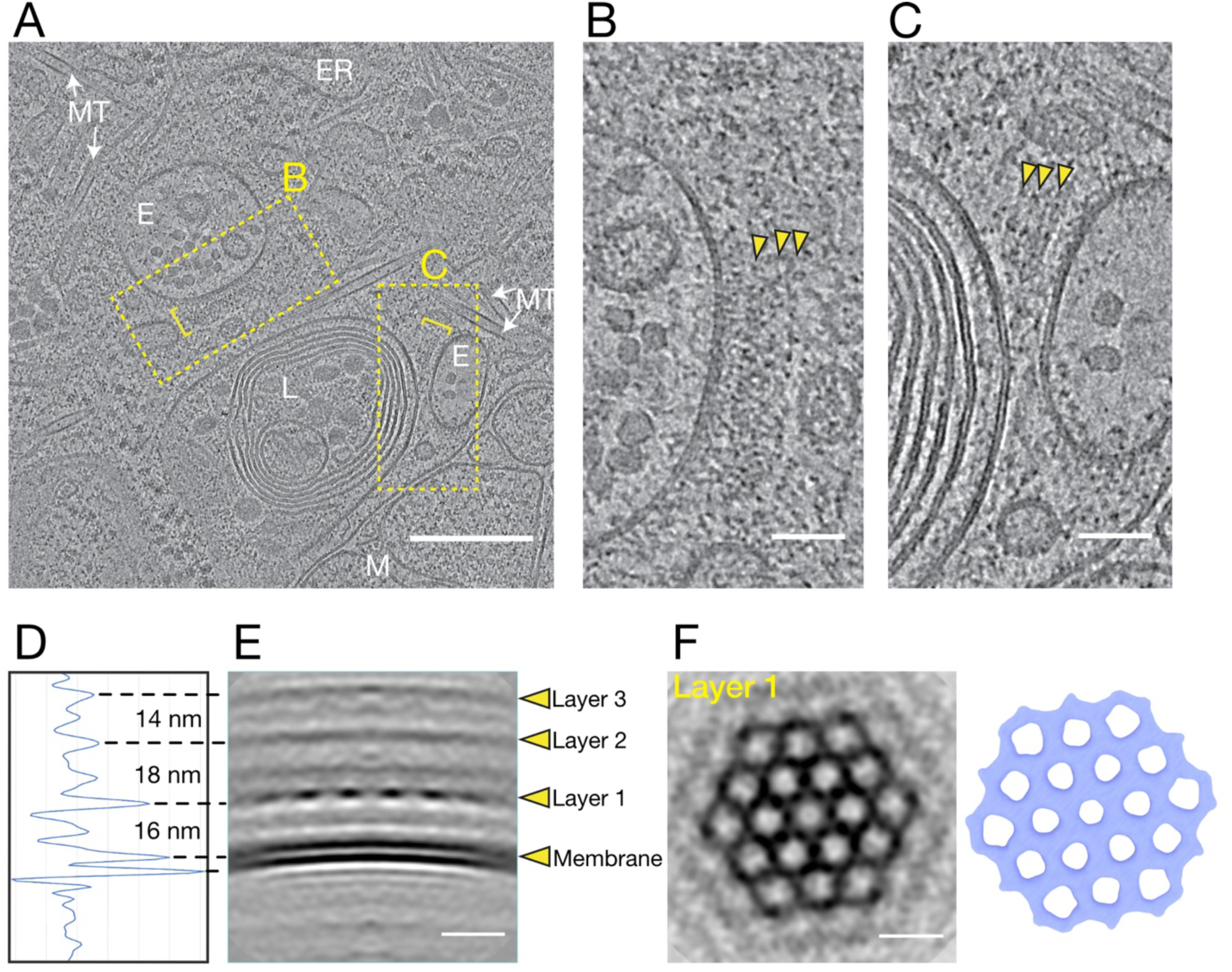
Clathrin assembles as a hexagonal lattice on endosomes. (A) A representative cryo-electron tomogram of endosomes with putative HRS-clathrin coat patches, indicated by yellow brackets. Areas magnified in panels b and c are indicated by yellow rectangles. White labels indicate some identified cellular components: E=endosomes, ER=endoplasmic reticulum, L=lysosomes, M=mitochondria, MT=microtubules. Scale bar is 250 nm. (B-C) Close-ups of protein coats on endosome membranes in tomogram from panel (A). Arrows point to the multiple layers within the coats. Scale bars are 50 nm. (D-F) Subtomogram averaging. (D-E) A density profile and side view of the membrane and protein layers with distances between the layers indicated. Scale bar is 20 nm. (F) A top view of layer 1 (perpendicular to view in e), shown as a slice through the volume and as an isosurface (blue), revealing the hexagonal lattice structure. Scale bar is 40 nm.

We next used subtomogram averaging to further analyze the organization of the coat (Figure S8A-D). This approach revealed a three-layer arrangement of the coat structure (Figure 4D-E). The first layer was positioned 16 nm from the endosomal membrane and showed a periodical pattern. Subsequent layers were 18 nm and 14 nm above the previous protein layers (Figure 4D). Distances of these protein layers from the membrane were strikingly similar to the height of HRS-clathrin assemblies measured by HS-AFM (Figure 2E and 3E). The periodic pattern of the first layer corresponded to a hexagonal array with a lattice constant (vertex-to-vertex length) of 24-26 nm (Figures 4F and S8E), while the same analysis of the second and the third layers did not reveal periodical arrangements. The dimensions of the lattice in the first protein layer are typical for flat clathrin lattices (Sochacki et al., 2021; Fotin et al., 2004; Kravčenko et al., 2024), indicating that this layer corresponds to clathrin predominantly polymerized into a hexagonal lattice.

Our *in vitro* reconstitution data shows that HRS readily assembles large, multi-layered assemblies of clathrin on PI(3)P-rich membranes. The appearance and dimensions of these reconstituted HRS-clathrin coats are highly similar to the multilayered and hexagonal architecture of clathrin coats found on endosomes in cells. We next asked what regulates the formation of these multilayered HRS-clathrin domains on endosomes and focused on the properties of the underlying membrane.

### Cholesterol enhances HRS and clathrin recruitment on endosomes in cells

Cholesterol is prominent at endosomes and other organelles where clathrin assembles (Van Meer et al., 2008; Kim and Burd, 2023; Bigay and Antonny, 2012). Therefore, we decided to test its role in clathrin assembly on endosomes. HRS forms distinct domains on endosomes (Raiborg et al., 2002, 2006) and we decided to test if these HRS domains are rich in cholesterol. We stained cholesterol in the mScarlet-I3-HRS cells with Filipin and observed colocalization of HRS with cholesterol (Figure 5A). We then tested if cholesterol levels affect the recruitment of HRS on endosomes. To this end, we used HeLa cells knock-out for the *NPC1* gene (Vacca et al., 2019), which encodes a lysosomal cholesterol transporter whose depletion elevates cholesterol in endosomal compartments (Pentchev et al., 1985; Liscum et al., 1989) (Figure 5B), and immuno-stained them against endogenous HRS. Remarkably, NPC1-KO cells displayed an increase in endosome-bound HRS-clathrin compared to control cells (Figure 5C-D), as seen from the significant increase in both HRS puncta size (Figure 5E) and clathrin fluorescence intensity (Figure 5F). We confirmed that HRS was still localized to endosomes in NPC1-KO cells, as endogenous HRS predominantly colocalized with EEA1, a marker of early endosomes, but not with LAMP1, a marker for lysosomes (Figure S9A-B). We also analyzed the HRS localization in cells treated with U18666A drug, an inhibitor of NPC1 (Liscum and Faust, 1989). Similar to NPC1-KO cells, U18666A-treated cells showed increased HRS signal in the endosomes (Figure S9C), while no change in HRS localization was observed (Figure S9D-E).

**Figure 5.**
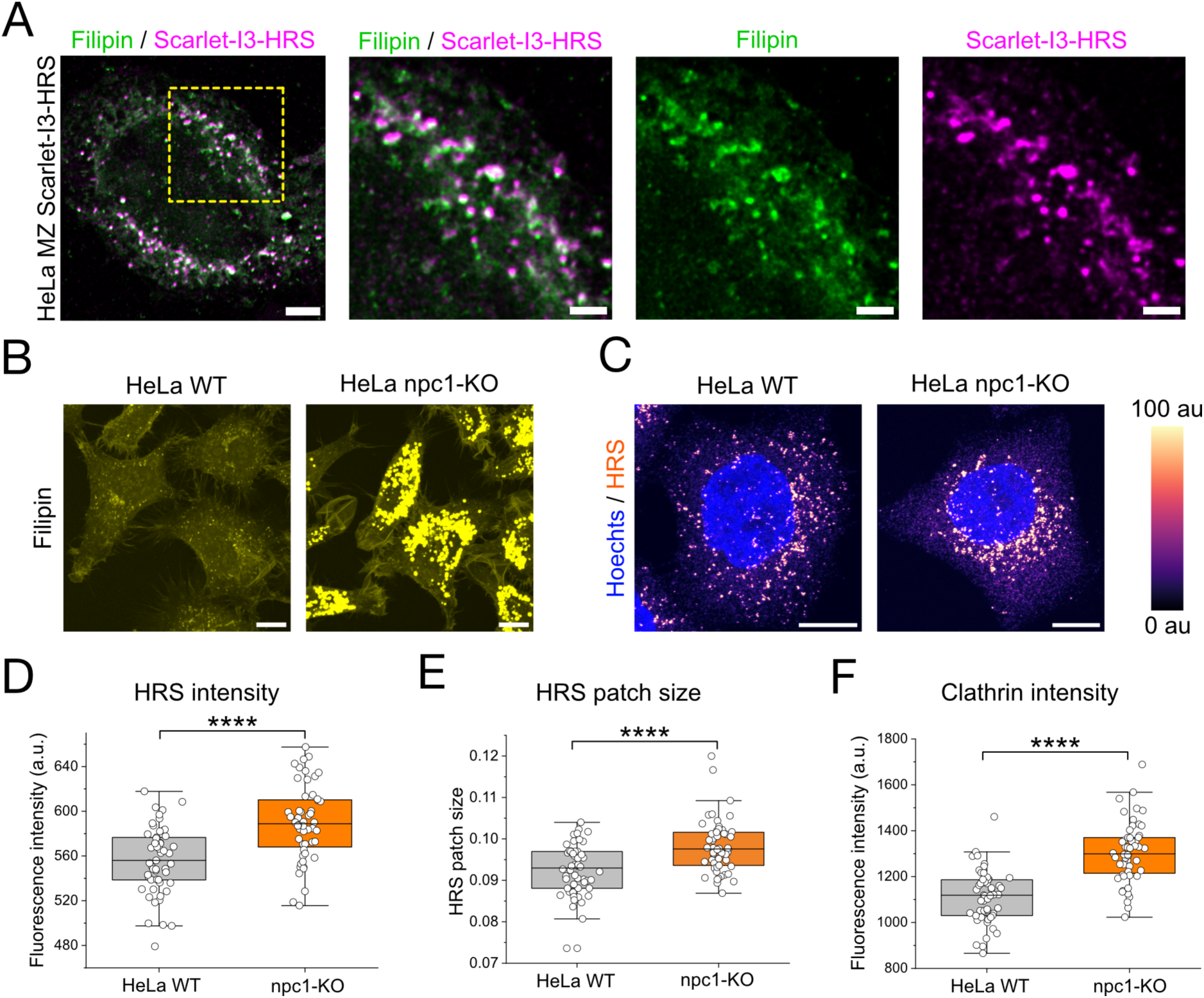
Cholesterol promotes endosomal localization of HRS in cells. (A) Representative maximum projection images from confocal fluorescence microscopy stack images of mScarlet-I3-HRS (magenta) CRISPR knock-in cell lines with filipin staining (green). The scale bar is 20 µm for larger images and 2 µm for zoom-in images. The experiment was repeated three times with similar results. (B) Filipin staining in HeLa wild-type cells and NPC1 knockout cells showing enrichment of cholesterol at endo-lysosomal membranes. Scale bars are 10 µm. (C) Immunofluorescence images of HRS in HeLa wild-type cells and NPC1 knockout cells. HRS staining is colored with a fluorescence intensity heatmap and nuclei (Hoechts) in blue. Scale bars are 10 µm. (D-F) High content image analysis of HRS mean intensity in cells (D), HRS mean patch size in cells (E), and clathrin mean intensity in HRS patches in cells (F). Each data point represents the mean of the image with median values, 25% and 75% percentiles, and the range of data indicated. Statistical testing: Kruskal-Wallis test, **** *p* < 0.0001.

We concluded from these data that HRS binds to endosomal membranes in correlation with their cholesterol content. To further understand the role of cholesterol in assembling HRS-clathrin coats, we tested its role in our *in vitro* reconstitution assay.

### Cholesterol promotes the assembly of the HRS-clathrin coat

We tested the recruitment of HRS on GUVs containing 1% PI(3)P and varying cholesterol concentrations: 0%, 15%, and 30% cholesterol (Lipid mixes 3, 1, and 4, respectively, Table S2). Cholesterol enhanced the recruitment of HRS on membranes (Figure 6A,C) and correspondingly, more clathrin was recruited on HRS-positive GUVs containing increasing amounts of cholesterol (Figure 6B,D). Interestingly, with 0% and 15% cholesterol, HRS was recruited on the membrane as small patches, whereas on membranes with 30% cholesterol, HRS either formed larger patches or covered the whole GUV membrane. We wondered if cholesterol could also promote the wetting of SLB membranes by HRS condensates. To test this, we prepared SLBs with varying cholesterol concentrations as above (0%, 15%, or 30% cholesterol) and tested the membrane wetting by using 2 µM fluorescently labeled AF488-HRS (Figure 6E). We observed a significant increase in membrane wetting with higher cholesterol concentrations (Figure 6F).

**Figure 6.**
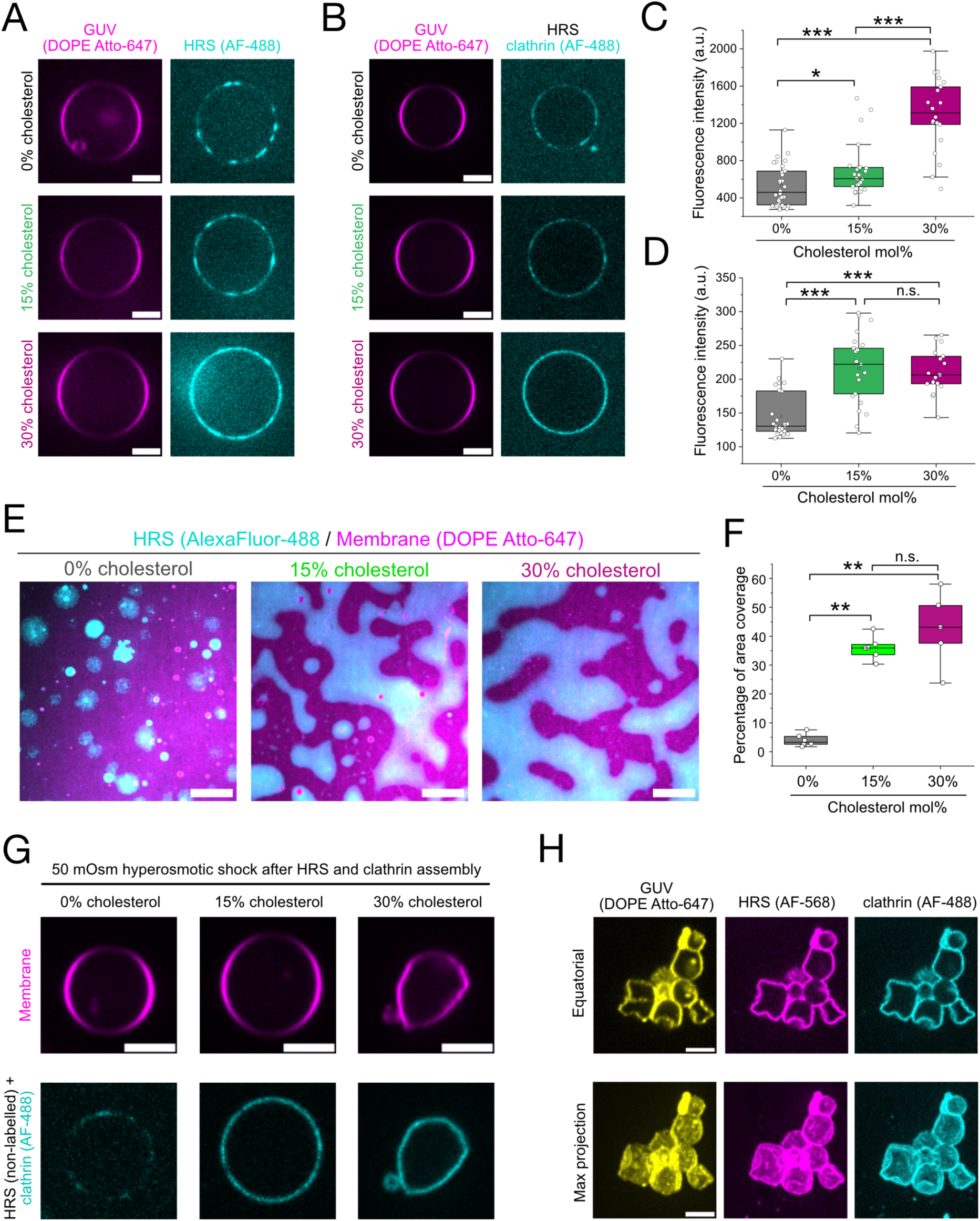
Cholesterol enhances the recruitment of HRS onto the membrane *in vitro*. (A) Representative fluorescence microscopy images of PI(3)P-rich GUVs (magenta) with indicated cholesterol concentrations and 500 nM labeled HRS (cyan). Scale bars are 5 µm. (B) Representative fluorescence microscopy images of PI(3)P-rich GUVs (magenta) with indicated cholesterol concentrations and 500 nM nonlabelled HRS and 200 nM clathrin (20% labeled, cyan). Scale bars are 5 µm. (C) The analysis of fluorescence intensity of labeled HRS on GUV membrane (from data in panel a) with different cholesterol concentrations. Each data point represents an individual GUV. The experiment was repeated three times (N=3). (D) The analysis of fluorescence intensity of labeled clathrin on GUV membrane with different cholesterol concentrations. Each data point represents an individual GUV. The experiment was repeated three times (N=3). (E) Representative fluorescence microscopy images of 2 µM labeled HRS (cyan) on PI3P-rich SLBs (magenta) with varying cholesterol concentrations. N=5 experiments for 0% and 30% cholesterol samples, and N=6 for 15% cholesterol samples. The scale bar is 50 µm. (F) The percentage labeled HRS-covered SLBs with different cholesterol concentrations. Each data point represents a mean coverage value in one sample. N=5 experiments for 0% and 30% cholesterol samples, and N=6 for 15% cholesterol samples. (G) Representative fluorescence microscopy images of GUVs first incubated with HRS and clathrin and subsequently brought into +50 mOsm osmolarity (hyperosmotic shock). Scale bars are 5 µm. The experiment was repeated three times (N=3) with similar results. (H) A representative example of flattened GUV membranes containing 30% cholesterol and incubated with 500 nM HRS and 200 nM clathrin before 50 mOsm osmotic shocks. Equatorial plane images and maximum projection images are shown. Scale bars are 5 µm. In panels (C), (D), and (F), each data point represents an individual measurement with median values, 25% and 75% percentiles, and the range of data indicated. Experiments were repeated three time (N=3). Statistical testing: Kruskal-Wallis test, * *p* < 0.05, ** *p* < 0.01, *** *p* < 0.001 **** *p* < 0.0001.

One of the key features of clathrin coats, especially when related to membrane trafficking, is their ability to bend the membrane. In our reconstitution samples, we observed both flat clathrin patches as well as regions where clathrin formed cup-shaped structures (Figures 1F-H and 3B-D). Moreover, we did not observe membrane deformation upon HRS-clathrin coat assembly on GUV membranes (Figure 6B), suggesting that the physiological assembly of HRS and clathrin may be flat. GUV membranes cannot be remodeled when membrane tension is too high, but lowering the tension through osmotic shocks promotes membrane remodeling by clathrin coats (Saleem et al., 2015). To test the membrane remodeling properties of HRS-clathrin assemblies, we coated GUVs with HRS and clathrin and then lowered the membrane tension with hyperosmotic shocks. HRS and clathrin reconstitution on GUVs containing 0% or 15% cholesterol and subjected to osmotic shock did not change shape (Figure 6G). In contrast, hyperosmotic shocks induced GUV flattening when HRS and clathrin were reconstituted on GUVs containing 30% cholesterol (Figure 6G). Such flattening, which appeared as facets on the GUVs (Figure 6H) and gives them the appearance of a broken line to the GUV contour, was previously observed with protein assemblies that preferred low curvature areas, such as Snf7 (Chiaruttini et al., 2015; Souza et al., 2025). Importantly, hyperosmotic shock did not lead to membrane deformations of GUVs without proteins or those incubated only with HRS (Figure S10A-B). Our experiments indicate that the HRS and clathrin indeed assemble as a flat coat and, under low membrane tension, induce the flattening of membranes (Figure 6G-H). This is in striking contrast with endocytic clathrin adaptor AP180 and clathrin that yielded prominent outward-directed budding assemblies upon hyperosmotic shocks (Saleem et al., 2015).

### Cholesterol clusters and diffuse slower under the HRS-clathrin coat

The finding that cholesterol enhances HRS wetting on the membrane led us to speculate that HRS could promote the formation of cholesterol-rich domains. This hypothesis could be assessed with a fluorescence quenching assay, which relies on the property of the TopFluor probe to self-quench at high density due to a fluorescence resonance energy transfer (FRET) between neighboring probe molecules (Figure S11A)(Zhao and Lappalainen, 2012). We prepared large unilamellar vesicles (LUVs) with 1% PI(3)P and 30% cholesterol supplemented with 0.1% TopFluor-cholesterol (Lipid mix 6, Table 2) and measured the fluorescence intensity of these vesicles with increasing concentrations of HRS in a spectrophotometer. TopFluor fluorescence decreased as a function of HRS concentration (Figure S11B), supporting the hypothesis that HRS clusters cholesterol. We then used SLBs labeled with 1% TopFluor-cholesterol (Lipid mix 5, Table S2) and tested if cholesterol gets enriched under HRS and clathrin. While it may look incoherent that HRS-dependent clustering of TopFluor-Cholesterol leads to global fluorescence quenching at the same time as fluorescence increase under the HRS domains, the lateral segregation of cholesterol under HRS domains could lead to apparition of fluorescent domains while promoting quenching because of the increased cholesterol density in HRS domains. Cholesterol was enriched under protein domains formed by both HRS condensates alone as well as HRS condensates incubated with clathrin (Figure S11C), supporting our hypothesis that cholesterol-rich domains form under HRS and HRS-clathrin patches.

The clustering of cholesterol underneath the HRS-clathrin coat should be associated with reduced diffusion, which can be measured by FRAP (Kefauver et al., 2024). To test this, we incubated membranes containing 1% TopFluor-cholesterol (Lipid mix 5, Table S2) with 500 nM HRS and 200 nM clathrin and photobleached small regions of membrane under the HRS condensates. Compared to the control without proteins, TopFluor-cholesterol showed a reduction of diffusion in samples incubated with HRS, and even more reduced diffusion when these samples were incubated also with clathrin (Figure S11D-F). These experiments indicate that HRS-clathrin coats decrease membrane fluidity and form cholesterol-rich, stable membrane domains, suggesting that lipid phase separation could be coupled to HRS-clathrin assemblies.

### HRS domains on cholesterol-rich membrane cluster cargo proteins

The main function of ESCRT-0 is to sort ubiquitylated cargoes into ILVs (Raiborg et al., 2002; Migliano et al., 2023; Banjade et al., 2022). We thus wondered whether HRS-clathrin could promote cargo clustering in our *in vitro* assay. To test this, we used the VAMP2 protein, a transmembrane cargo model for ESCRT-0 in previous studies (Takahashi et al., 2015). We purified a recombinantly expressed VAMP2 chimera with four ubiquitins at its N-terminus and sfGFP at its C-terminus and reconstituted it into the GUV membrane (see Methods and Figure 7A). To test the effect of cholesterol on cargo clustering, we prepared cargo-containing membranes either with or without cholesterol (Lipid mixes 3 and 4, respectively, Table S2). Without HRS, Ub-VAMP2-GFP showed mostly homogenous distribution in GUVs, with some domains of higher intensity, in both tested lipid compositions (Figure 7B). When we incubated these cargo-containing GUVs with HRS, cargo clusters appeared to be larger and more stable in membranes containing 30% cholesterol compared to those without cholesterol (Figure 7C, Movie S6-7), indicating that cholesterol domains promote HRS-clathrin cargo clustering. Our data thus reveal that HRS-clathrin assemblies are sufficient to sort ubiquitinylated cargoes *in vitro*, and that this clustering is enhanced by cholesterol.

**Figure 7.**
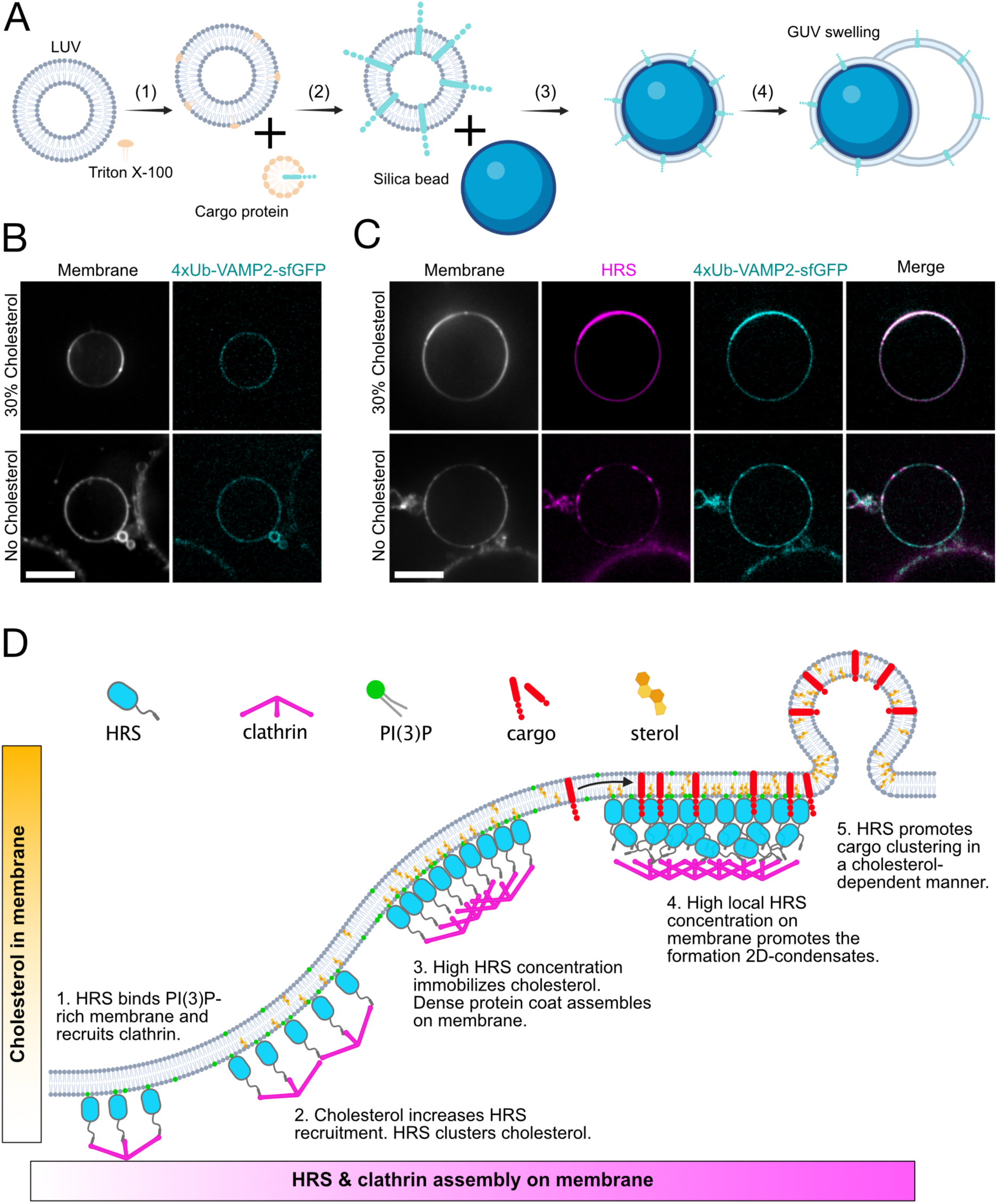
HRS clusters ubiquitylated cargoes in a cholesterol-dependent manner. (A) A schematic illustration of reconstitution (see Materials and Methods for details) of 4xubiquitin-VAMP2-sfGFP cargo protein into GUV membrane (B) Representative fluorescence microscopy images of 4xubiquitin-VAMP2-sfGFP (cyan) reconstituted on GUV membrane (grey) containing either 1% PI(3)P/30% cholesterol or 1% PI(3)P/no cholesterol before the addition of HRS. The scale bar is 5 µm. (C) Representative fluorescence microscopy images of 4xubiquitin-VAMP2-sfGFP (cyan) reconstituted on GUV membrane (grey) containing either 1% PI(3)P/30% cholesterol or 1% PI(3)P/no cholesterol after addition of 500 nM mScarlet-I3-HRS (magenta). The scale bar is 5 µm. The experiment was repeated four times (N=4) with similar results. (D) A model of HRS and clathrin assembly on PI(3)P-rich membrane in a cholesterol-dependent manner. HRS binds to PI(3)P through its FYVE domain (1), and clusters cholesterol, which leads to increased HRS recruitment on the membrane (2) and finally to the formation of a stable, cholesterol-rich membrane domain under dense HRS clathrin lattice (3). High local protein concentration leads to the formation of two-dimensional HRS condensates. High-density HRS lattice and stable cholesterol domain under the protein promote cargo sorting under the HRS-clathrin lattice (4). All HRS populations recruit clathrin. Ubiquitinated cargoes get clustered under the HRS-clathrin lattice, and cholesterol stabilizes these HRS-cargo clusters (5).

## Discussion

While functions of flat clathrin lattices in cellular processes have gathered increased attention during the past years (Vassilopoulos and Montagnac, 2024), it remains unknown how these structures are assembled in the first place. Here, we reconstituted endosomal, ESCRT-0-driven clathrin coats with minimal components to gain a comprehensive understanding of their assembly mechanism and structural features. We reveal that HRS directly interacts with the membrane, initially forming a low-density protein coat which can also phase-separate into a dense, two-dimensional protein film on the membrane. Clathrin assembles on HRS-rich membrane as a dense coat which has either a flat or cup-shape architecture depending on the nature of the condensate it binds to (two-dimensional films versus three-dimensional droplets). Our experiments with GUVs show that the dense HRS-clathrin coat flattens low tension membranes, indicating that the flat architecture is a dominant feature of the HRS-clathrin coat. This is in striking contrast with the reconstitution experiments performed with CME-linked clathrin adaptors, which promote membrane bending (Saleem et al., 2015; Dannhauser and Ungewickell, 2012; Kovtun et al., 2020), highlighting the unique features of the HRS-clathrin coat among clathrin assemblies.

Cryo-electron tomography data revealed flat HRS-clathrin protein coats in cells. These coats appeared to be composed of three layers with an architecture different from clathrin structures found on the plasma membrane (Heuser, 1980; Avinoam et al., 2015; Vassilopoulos and Montagnac, 2024) and consistent with previously published conventional cellular electron microscopy data (Sachse et al., 2002; Wenzel et al., 2018; Raiborg et al., 2001a, 2002). This layering of clathrin structures is also seen in our in vitro samples. Notably, two-dimensional HRS condensates and HRS-clathrin coats on SLBs had a thickness of 10 nm and 20-30 nm, respectively, which are in the range of the measured distance between the membrane and the clathrin lattice on endosomes. Besides the fact that the in vivo and in vitro structures are multi-layered with similar thicknesses, the layer closer to the membrane contains clear flat clathrin lattices both in vivo and in vitro. Sub-tomogram averaging revealed a hexagonal lattice 16 nm away from the endosome membrane, which resembled other clathrin assemblies imaged in cells (Sochacki et al., 2021; Heuser, 1980; Fotin et al., 2004; Kravčenko et al., 2024; Tagiltsev et al., 2021). Supporting a two-dimensional condensate of HRS forming the ESCRT-0 structure, we did not observe a distinct protein structure between the clathrin layer and the membrane, where HRS is expected to localize. This observation could be explained by the intrinsically disordered nature of HRS, which will result in “averaging out” relative to the clathrin lattice. We observed two additional protein layers on top of the hexagonal clathrin lattice on endosomes. These layers did not show apparent symmetry and the identity of these additional layers remains to be solved. We speculate that the three layers of this unique coat could be stacks of two-dimensional condensates separated by different types of clathrin assemblies. The flat nature of the HRS-clathrin coat may be due to the higher rigidity, associated with the loss of diffusion, of the two-dimensional HRS condensate that could force clathrin to adapt a flat layer.

Protein phase separation through IDRs is a phenomenon that has recently been shown to play an important role in membrane remodeling and membrane repair (Ravindran and Michnick, 2024; Yuan et al., 2021; Mangiarotti et al., 2023; Bussi et al., 2023; Schiano Lomoriello et al., 2022). In clathrin-mediated endocytosis, Eps15 and its budding yeast homolog Ede1 were shown to undergo liquid condensation (Kozak and Kaksonen, 2022; Day et al., 2021). However, to our best knowledge, HRS is the first protein directly interacting with clathrin that undergoes condensation. Thus, HRS and clathrin are unique in combining a liquid-like protein condensate and a well-structured protein lattice in the same protein assembly and give a plausible explanation why clathrin coats on endosomes have distinct, multilayered architecture compared to clathrin structures on the plasma membrane and on other organelles. In addition, HRS phase separation may be needed to circumvent the ESCRT-0 domains in delimited regions of endosomes, as without phase separation, the HRS-clathrin coat could grow indefinitely because it is flat.

We show that when HRS and clathrin form a protein coat on the membrane, they change the membrane composition by enriching cholesterol under the protein coat. Here, cholesterol shows lower diffusion, which suggests that HRS and clathrin can promote the formation of cholesterol-rich membrane domains. Since HRS does not contain cholesterol-binding domains or motifs, cholesterol enhances HRS recruitment via an indirect mechanism. Cholesterol was shown to increase the phosphoinositide-dependent membrane interaction with other proteins by clustering phosphoinositides (Lolicato et al., 2022). Moreover, the interaction of protein condensates with membranes is coupled with lipid phase separation (Wang et al., 2023; Snead and Gladfelter, 2019). We speculate that HRS combines these mechanisms to promote cholesterol clustering and membrane domain formation. HRS-clathrin coats form distinct protein domains on endosomes (Figure 4A-C and S9e), which were also reported by other studies (Raiborg et al., 2002, 2006; Wenzel et al., 2018; Norris et al., 2017). How these protein domains are assembled in the first place and how their size is regulated have remained unanswered questions. We propose that the feedback loop between HRS recruitment and the formation of cholesterol-rich membrane domains (Figure 7D) facilitates the rise of HRS-positive microdomains on endosomes. Supporting this, the depletion of CD63, a tetraspanin protein that sorts cholesterol to ILVs on endosome membranes (Palmulli et al., 2024), results in an unusual abundance of multilayered clathrin coats on multivesicular endosomes (van Niel et al., 2011).

Niemann-Pick C (NPC) 1 and NPC2 proteins at lysosomes are key regulators of cholesterol homeostasis in cells (Meng et al., 2020; Ikonen and Zhou, 2021; Ikonen, 2008). Our observation that cholesterol depletion increases the HRS localization at endosomes highlights that cholesterol homeostasis is an important factor regulating ESCRT function and, more globally, the endosomal activity in cells. Sterol depletion at the plasma membrane inhibits membrane bending and vesicle invagination in the CME (Anderson et al., 2021; Subtil et al., 1999; Rodal et al., 1999; Pichler and Riezman, 2004), which highlights the importance of sterol homeostasis in cells. To our knowledge, the effect of sterols on clathrin adaptor recruitment and clathrin assembly on the plasma membrane has not been studied thoroughly. However, it would not be surprising if cholesterol affects clathrin adaptor recruitment on cellular membranes in a similar fashion to HRS. It would thus be interesting to test if clathrin adaptor proteins linked to endocytosis, long-lived plasma membrane clathrin plaques, or reticular adhesions are regulated by sterols.

In conclusion, we reveal the unique mechanism of how endosomal protein microdomains of ESCRT-0 and clathrin are formed. Clathrin is assembled as a hexagonal lattice on HRS, which forms two-dimensional condensates on membranes. The interplay between the cholesterol-rich membranes and HRS condensates facilitates the protein recruitment on the membrane and formation of a cholesterol-rich diffusion barrier in the membrane. This mechanism also provides a platform for cargo clustering under the HRS-clathrin microdomain. Future studies will reveal how cargo proteins are transported from the HRS-clathrin domain to the intraluminal vesicle.

## Methods

### Plasmids and protein purification

All lipids used in this study are commercially available at Avanti Polar Lipids and are listed with their respective catalog numbers in Table S1. The vector pCoofy17 was a gift from Sabina Suppmann (Addgene plasmid # 43991)(Scholz et al., 2013). A plasmid pCS2 HRS-RFP was a gift from Edward de Robertis (Addgene plasmid # 29685). pCoofy17 and HRS gene were linearized with PCR and subsequently assembled with sequence and ligase independent cloning approach as described in (Scholz et al., 2013). Plasmids for expression of mScarlet-I3-HRS [pET51(+)-mScarlet-I3-HRS) and 4xUbiquitin-VAMP2-sfGFP [pET28(+)-4xUb-VAMP2-sfGFP] in bacteria, and mScarlet-I3-HRS overexpression in mammalian cells [pCDNA3.1(Zeo+)-mScarlet-I3-HRS] were synthesized and cloned into the destination vector by Gene Universal Inc.

All recombinant protein expressions were done in BL21 pLysS bacteria either in standard LB media with 0.5 mM IPTG induction or in Autoinduction LB media (# AIMLB0210, Formedium). All proteins were expressed overnight at 20°C. For HRS protein purification, cells were sonicated in HRS lysis buffer (50 mM Tris-HCl, pH 8.0, 500 mM NaCl, 5 mM 2-mercaptoethanol, 300 mM L-arginine) supplemented with PMSF and c0mplete protease inhibitor cocktail (#11697498001, Roche). Soluble protein was then bound to Talon cobalt resin, washed three times with HRS lysis buffer, and then three times with HRS wash buffer (50 mM Tris-HCl, pH 8.0, 500 mM NaCl, 5 mM imidazole, 5 mM 2-mercaptoethanol, 300 mM L-arginine). HRS was cleaved from Sumo3 tag on resin with SenP2 protease in elution buffer (20 mM Tris-HCl, pH 8.0, 500 mM NaCl, 5 mM 2-mercaptoethanol, 300 mM L-arginine pH 7.4). The final protein was cleaned with a Superdex-200 size exclusion column (Cytiva) equilibrated with the elution buffer using Äkta Pure system (Cytiva). A 5% final concentration of glycerol was added to the protein before snap freezing with liquid nitrogen and storage at −80°C.

To purify VAMP2, cells were lysed in VAMP2 lysis buffer (20 mM Tris-HCl, pH 8.0, 300 mM NaCl, 0.1% Tween-20, 1% Triton X-100, 30 mM imidazole) supplemented with PMSF and c0mplete protease inhibitor cocktail (#11697498001, Roche) using sonicator. Cleared protein lysate was then bound to HisTrap 5 ml HP column and washed with 20 column volumes of VAMP lysis buffer. The protein was eluted with 300 mM imidazole in VAMP lysis buffer and subsequently cleaned with Superdex-200 size exclusion column (Cytiva) equilibrated with the VAMP2 elution buffer (Lysis buffer without imidazole) using Äkta Pure system (Cytiva). A 10% final concentration of glycerol was added to the protein before snap freezing with liquid nitrogen and storage at −80°C. Buffer (Lysis buffer without imidazole) using Äkta Pure system (Cytiva). A 10% final concentration of glycerol was added to the protein before snap freezing with liquid nitrogen and storage at −80°C.

To purify mScarlet-I3-HRS, cells were lysed in HRS lysis buffer supplemented with PMSF and a c0mplete protease inhibitor cocktail (#11697498001, Roche) using Emulsiflex. Protein was then bound to Step-Tactin XT 4Flow resin (Iba Life Sciences). The resin was then washed with HRS lysis buffer, and the protein was eluted with 5 mM biotin in HRS lysis buffer. The final protein was cleaned with Superdex-200 size exclusion column (Cytiva) equilibrated with the HRS elution buffer using Äkta Pure system (Cytiva). A 5% final concentration of glycerol was added to the protein before snap freezing with liquid nitrogen and storage at −80°C.

Clathrin from fresh bovine brains was purified with a standard protocol as described in (Ahle and Ungewickell, 1986). Purified clathrin and HRS were labeled with either AlexaFluor-488 or AlexaFluor-568 conjugated C5 maleimide (# A10254 and # A20341, respectively, Thermo Fisher Scientific Scientific) according to the manufacturer’s protocol.

### Reconstitution on giant unilamellar vesicles (GUV)

GUVs were prepared with the gel-assisted swelling method as described in (Weinberger et al., 2013) or with silica-bead assisted swelling method. Shortly, PVA (MW 145000, # 8.14894.0101, Sigma-Aldrich) was dissolved at 5% (w/v) in 280 mM sucrose solution. Glass coverslips were cleaned by sonicating them sequentially in water, ethanol, acetone, 1 M KOH and finally in water. A thin layer of dissolved PVA was spread on glass and dried in a 50 °C oven for 30 min. Lipids were mixed in chloroform to the desired composition (Table S2) at 1 mg/ml final concentration, and 10 µl of this lipid mixture was spread on PVA gel. Chloroform was then evaporated for 30 min at a 30°C vacuum oven. 500 µl of swelling buffer (20 mM HEPES, pH 7.4, 50 mM NaCl, osmolarity was adjusted with sucrose) was then added on dried lipids, and GUVs were swelled at room temperature for 30 min.

To prepare GUVs with a silica bead method, we mixed lipids in chloroform at the desired lipid composition (Table S2) at 0.5 mg/ml concentration and evaporated chloroform first under argon gas and then in a vacuum for 60 minutes. Lipids were then hydrated with 5 mM Hepes, pH 7.4 buffer, and subsequently mixed with silica beads (# 904384, Sigma Aldrich). Lipids and silica beads were dried in a vacuum for 30 minutes and then prehydrated with 1 M trehalose solution. Prehydrated, membrane-coated silica beads were then transferred in HRS-clathrin reaction buffer (20mM HEPES pH 7.2, 125 mM KAc, 1 mM MgAc).

Protein binding reactions with GUVs were performed in HRS-clathrin reaction buffer (20 mM HEPES pH 7.2, 125 mM KAc, 1 mM MgAc). The osmolarity of the reaction buffer was adjusted to match the reaction buffer and GUV swelling buffer with sucrose, and osmolarity was always measured before experiments with Vapor Pressure Osmometer (Wescor). The osmolarity of the reaction buffer was increased with sucrose for hyperosmotic shock experiments. Labeled proteins were incubated with GUVs in a closed reaction chamber (Ibidi Sticky Slide IV, # 80608, Ibidi GmbH) on a cover glass passivated with either 2 mg/ml BSA in reaction buffer or with PEG-silane. Imaging was performed with a spinning disk confocal microscope (3i Intelligent Imaging Innovations) built on Nikon Eclipse C1 using a 100x 1.3 NA objective. The microscope was controlled with Slidebook 6.0.22 (3i Intelligent Imaging Innovations), and data analysis was performed using Fiji v1.54f. Final graphs were generated with OriginPro 2024b v10.1.5.132.

### Reconstitution on supported lipid bilayers (SLBs)

To prepare SLBs, we mixed lipids in chloroform and membrane-coated silica beads as described above. Membrane-coated silica beads were then hydrated on plasma-cleaned coverslips attached to Ibidi Sticky Slide IV (# 80608, Ibidi GmbH) in HRS-clathrin reaction buffer (20mM HEPES pH 7.2, 125 mM KAc, 1 mM MgAc). Supported bilayers were spread on glass for 15-30 minutes, and the remaining bare glass areas were passivated with 2 mg/ml BSA in reaction buffer for 60 minutes. After the membrane was washed five times with the reaction buffer, HRS protein was added into the sample chamber in the desired concentration in the reaction buffer. In clathrin assembly assays, HRS was incubated on the membrane for 60 minutes before the sample chamber was washed five times with a reaction buffer. To avoid unwanted assembly of clathrin cages in solution, all experiments were performed at physiological pH and salt concentrations. Additionally, we reduced the in-solution interaction between HRS and clathrin by sequentially introducing proteins onto supported lipid bilayers (SLBs) using microfluidic chambers. After incubating HRS for 60 minutes, we washed the chambers five times with reaction buffer before adding clathrin.

The assembly of labeled HRS and clathrin and fluorescence recovery after photobleaching (FRAP) of assembled protein coats were recorded using an Olympus IX83 TIRF microscope equipped with ImageEM X2 EM-CCD camera (Hamamatsu), Olympus Uapo N 100x NA 1.49 objective. The microscope was controlled with Visiview v.4.4.0.11 software (Visitron Systems). Data was analyzed with Fiji v1.54f, and final graphs were generated with OriginPro 2024b v10.1.5.132.

### Protein diffusion measurements

Protein coat diffusion after the recruitment was measured by using the Fluorescence Recovery After Photobleaching (FRAP) approach using an Olympus IX83 TIRF microscope as above. SLBs were incubated with desired concentrations of labeled HRS, and the protein recruitment was followed over time until the fluorescence intensity plateaued. The remaining bulk protein was then washed out with five consecutive washes with the reaction buffer. The diffusion of membrane-bound protein was measured immediately by FRAP. A small region of protein on the membrane was photobleached, and the recovery of fluorescence was monitored by time-lapse imaging with 1-second intervals. Data was analyzed with Fiji v1.54f, and final graphs were generated with OriginPro 2024b v10.1.5.132.

### Platinum replica electron microscopy (PREM)

PREM samples were prepared as described above with the exception that instead of closed sample chambers, we prepared samples in open chambers, which were prepared by attaching a microscope coverslip on a round metal, homemade sample holder. After the assembly of HRS and clathrin, samples were washed five times with a reaction buffer and then fixed with 2% glutaraldehyde/2% formaldehyde solution for 30 minutes. Samples were stored in 2 % glutaraldehyde in reaction buffer for 16-40 hours until the PREM replicas were prepared. SLBs were then sequentially treated with 0.5% OsO_4_, 1% tannic acid, and 1% uranyl acetate before graded ethanol dehydration and hexamethyldisilane substitution (Sigma-Aldrich). Dried samples were rotary-shadowed with 2 nm of platinum and 6 nm of carbon using a high vacuum sputter coater (Leica Microsystems). The resultant platinum replica was floated off the glass with hydrofluoric acid (5%), washed several times with distilled water, and picked up on 200 mesh formvar/carbon-coated EM grids. Replicas on EM grids were mounted in a eucentric side-entry goniometer stage of either a transmission electron microscope operated at 80 kV (model CM120; Philips) equipped with a Morada digital camera (Olympus) or a transmission electron microscope running at 120 kV (Jeol) equipped with a Xarosa digital camera (EMSYS GmbH). Images were adjusted for brightness and contrast in Adobe Photoshop (Adobe) and presented in inverted contrast.

### Quantification of multilayered clathrin coats

After acquiring images, they were processed with ImageJ software to adjust brightness and contrast, ensuring that electron opacity is visible and quantifiable. To quantify the density of clathrin layers, line profiles were drawn across regions of interest on various clathrin layers in the PREM images. ImageJ was then used to analyze these line profiles, allowing for precise quantification of the gray values that correspond to electron opacity or intensity. The analysis differentiates between various clathrin layers based on their line profiles by identifying distinct maxima and background levels. For instance, monolayer and double clathrin layers are characterized by two distinct maxima in their line profiles, with gray values that exceed the background levels. A gray value above the background level serves as a criterion to confirm the presence of an additional clathrin layer.

### Fast Atomic force microscopy and Quantitative imaging

AFM images were acquired using a JPK NanoWizard Ultraspeed AFM (Bruker and JPK BioAFM) equipped with USC-F0.3-k0.3-10 cantilevers (Nanoworld) with a spring constant of 0.3 N/m and a resonance frequency of approximately 300 kHz. The Fast-AFM was operated in tapping mode, with the cantilever oscillating at a frequency close to 150 kHz.

Large unilamellar vesicles (LUVs) composed of DOPC:DOPS:DOPE:PI(3)P:Ch (29:20:20:1:30) were spread onto freshly cleaved mica surfaces to form supported lipid bilayers (SLBs) using buffer extension (10 mM HEPES, 2 mM MgCl₂, 10 mM CaCl₂, pH 7.4) at 37°C for 30 minutes. The samples were then gently rinsed three times with reaction buffer to remove excess non-bound lipids. SLBs were imaged before selecting and scanning the areas of interest (AOIs).

During imaging, HRS and clathrin samples were injected into the imaging chamber at final concentrations of 500 nM and 200 nM, respectively. First, the dynamics and mechanics of HRS were measured for at least 30 minutes using Fast-AFM imaging and quantitative Imaging (QI) mode, followed by clathrin injection and subsequent measurement. To minimize lateral frictional forces during clathrin injection and its interaction with HRS, QI mode was used. This mode provides quantitative nanomechanical data maps correlated with topography. A low force regime of 600–6800 pN was applied. The vertical length and forward velocity parameters were set to 100-250 nm and 50-120 µm/s depending on the topology height of the scanned area, respectively. Images were analyzed using JPKSPM Data Processing, Fiji, and WSxM software (Horcas et al., 2007).

### Cell lines

HeLa MZ wild type and NPC1 knockout cell lines were a kind gift from Stefania Vossio (ACCESS, University of Geneva) (Vacca et al., 2019). HeLa Kyoto cell line constitutively expressing mCherry-HRS was a kind gift from Harald Stenmark (Faculty of Medicine, University of Oslo). We maintained these cells in high-glucose DMEM + GlutaMax-I (# 61965-026, Thermo Fisher Scientific) supplemented with penicillin-streptomycin solution (# 15140122, Thermo Fisher Scientific), 10% fetal bovine serum (# 10270-106, Thermo Fisher Scientific) and 1 mM sodium pyruvate (# 11390070 Thermo Fisher Scientific) in 37°C and 5% CO_2_ cell culture incubator.

We generated the mScarlet-I3-HRS (Figure S6A) cell line by targeting the N-terminal part of the HRS genomic locus (ID 9146) with the guide RNA: 5’ –GGTGCCGCTGCCTCGCCCCA<u>TGG</u>-3’ [<u>P</u>rotospacer <u>A</u>djacent <u>M</u>otif <u>S</u>equence (PAM) is underlined] located at the start codon in the HRS coding sequence (PAM just before start codon). The designed guide RNA was purchased from Genescript. The guide RNA was designed using Benchling (https://www.benchling.com) and double-checked using the CRISPR-Cas9 guide RNA design checker from IDT (https://www.eu.idtdna.com/site/order/designtool/index) to analyze on-target and off-target score. For this guide RNA, no significant off-targets were detected. The Homology Repair (HR) primers (sense and anti-sense) were designed to be adjacent to the start codon from the genomic sequence and with a length of 55 nucleotides each. The primers used for HR templates were: Forward and HRS-Right-Homology-Arm sense: 5’-GCGCCCGCGGCGTCGGGTTTGGGCTGGAGGTCGCCATGGATAGCACCGAGGCCGT-3’; Reverse and HRS-Left-Homology-Arm anti-sense: 5’-AGACCACCTCCTCCTAGATCGTACCCCGCTCCGTCG CCGTGGAAGCTCGCAGAGGA-3’. The DNA sequence of the right and left homology arms was 35 nucleotides, and the sequence to hybridize with the fluorescent tag was 20 nucleotides. The primers were synthesized by Microsynth AG. The fluorescent tag mScarlet-I3 was synthesized *de novo* by Genescript. This fragment was cloned into a pUC57 plasmid using EcoRI and BamHI restriction enzymes. For the generation of mScarlet-I3, the amino acid sequence was obtained from FPbase (https://www.fpbase.org), and then, by using Genescript and Benchling algorithms, the optimized DNA codon sequence was obtained for best mammalian expression. To the 3’-terminus of mScarlet-I3, a long flexible linker (69 amino acids) was added to keep the maximum functionality of the tagged protein. This linker is designed to have multiple GS-rich regions, two times the ALFA epitope (5’-CCCAGCAGACTGGAAGAGGAAC TGCGGCGGAGACTGACAGAA -3’) for Western Blott detection of the inserted tag, and a TEV cleavage site (5’-GAGAACCTGTACTTTCA AGGCGCCGCTAAGTTC-3’) for downstream protein purification. The following double-strand DNA fragment was inserted into the pUC57 plasmid: mScarlet-I3-GS-2xALFA-GS-TEV-GS: 5’-ATGGATAGCACCGAGGCCGTGATCAAAGAGTTCATGCGGTTCAAGGTGCACATGGAAGGCAGCATGAACGGCCACGAGTTCGAGATCGAAGGCGAAGGCGAGGGCAGACCTTATGAGGGAACACAGACCGCCAAGCTGAAAGTGACCAAAGGCGGCCCTCTGCCTTTCAGCTGGGACATTCTGAGCCCTCAGTTTATGTACGGCAGCCGGGCCTTCATCAAGCACCCTGCCGATATTCCCGACTACTGGAAGCAGAGCTTCCCCGAGGGCTTCAAGTGGGAGAGAGTGATGATCTTCGAGGACGGCGGCACCGTGTCTGTGACCCAGGATACAAGCCTGGAAGATGGCACCCTGATCTACAAAGTGAAGCTGAGAGGCGGCAACTTCCCTCCTGATGGCCCCGTGATGCAGAAAAGAACCATGGGCTGGGAAGCCAGCACCGAGAGACTGTACCCTGAGGACGTGGTGCTGAAGGGCGACATCAAGATGGCCCTGAGACTGAAGGATGGCGGCAGATACCTGGCCGACTTCAAGACCACCTACAAGGCCAAGAAACCCGTGCAGATGCCAGGCGCCTTCAACATCGACCGGAAGCTGGATATCACCAGCCACAACGAGGACTACACCGTGGTGGAACAGTACGAGAGAAGCGTGGCCAGACACAGCACAGGTGGAAGCGGAGGATCTTCCTCAACTGGTGGTGGAGGTTCTAGTCCCAGCAGACTGGAAGAGGAACTGCGGCGGAGACTGACAGAAGGCGGCGGAGGATCTTCTGGCCCTTCTGGATCTAGCAGCCCCTCCAGGCTGGAGGAAGAACTGAGAAGAAGGCTGACCGAGGAGAACCTGTACTTTCAAGGCGCCGCTAAGTTCAGCTCTGGTGGAGGAGGATCTAGC -3’.

For the generation of the double-strand DNA Donor Template, we produced PCR cassettes using the plasmid generated before together with the HR Right-Left primers to amplify the tag with the 35 nucleotide-Homology Arms. For the PCR amplification, the PrimeSTAR Max DNA Polymerase kit (#R045A) from Takara was used. For the purification of the PCR product, the QIAquick PCR and gel Cleanup kit (28506) from Qiagen was used. HeLa MZ cells were seeded into a 24-well plate at 1×10^5^ cells/mL without antibiotics one day before transfection. We used the Lipofectamine CRISPRMAX Cas9 (CMAX00008) as a Transfection Reagent from Thermofisher, and we followed the manufacturer’s indications for the transfection. Together with the transfection system, we used the purified GenCrispr NLS-Cas9-EGFP Nuclease (Z03467) from Genescript, the guide RNA designed to target HRS, and the HR PCR cassettes generated as a template (500 ng/well). Just before adding the Cas9/gRNA complex, the media was replaced with new media without antibiotics, and Alt-R HDR Enhancer V2 (10007921) from Integrated DNA Technologies to increase the chances of Homology-Directed Repair events after the DNA double-strand break by the Cas9 nuclease. One day later, the media was replaced for new media with antibiotics. To obtain single-cell clones of mScarlet-I3-HRS Knock-In, cells were subjected to FACS, and single cells were deposited in a well of a 96-well plate. For the validation of the cell lines, clones were genotyped via Western Blot using anti-HRS and anti-ALFA antibodies (see Table S4), fluorescent microscopy, and PCR using the following primers: HRS Forward 5’-GTTCTTAGGGCTCATTGTTCCA-3’ targeting the 5’-UTR, and therefore, before the start codon; and HRS Reverse 5’-AGTTCACTCTGTGGAAGGAACG-3’ targeting the HRS coding sequence after the start codon. For the genomic DNA extraction, the QIAamp DNA Mini kit (51304) from Qiagen was used. Final validation was performed, followed by sequencing PCR amplified regions to check the correct insertion of the tag using Microsynth AG sequencing service.

Western blot analysis revealed that one of the clonal cell lines was heterozygous for HRS-mScarlet-I3, while the other one was homozygous for HRS-mScarlet-I3. We noted that tagging of HRS decreased its expression levels (Figure S6B), and therefore used a heterozygous cell line, as it has total HRS expression levels closer to the ones of non-modified cells.

### Cryo-CLEM and cryo-ET

HeLa MZ cells with endogenous HRS tagged with Scarlet-I3-HRS and HeLa Kyoto cells stably expressing HRS-mCherry (Wenzel et al., 2018; Migliano et al., 2023) were grown on 200 mesh gold EM grids with a holey carbon film R2/2 (Quantifoil). Cells were incubated with 50 ng/ml EGF-Alexa 647. After allowing internalization of the cargo for 15 minutes at 37°C, the remaining EGF-Alexa647 was washed away with cell culture media. The EM grids were backside-blotted for 12 seconds using Whatman No.1 filter paper and vitrified using a manual plunger with a cryostat (Russo et al., 2016) between 7 and 12 minutes after the removal of EGF-Alexa647.

Thin lamellae of the cells were obtained by cryo-focused ion beam milling in an Aquilos 2 cryo-FIB-SEM (Thermo Fisher Scientific), equipped with an integrated fluorescence light microscope (iFLM). Grids were subjected to platinum coating for 1 minute and 30 seconds. Lamellae were prepared using standard semi-automated protocols (Wagner et al., 2020; Ganeva et al., 2023). Eucentricity, finding milling angles, stress relief cuts, and rough and medium milling steps were performed using the Cryo AutoTEM software in an automatic manner (Thermo Fisher Scientific). After assessing rough-milled lamella for signals of interest by iFLM, final polishing to a target thickness of 200-250 nm was performed manually. In some cases, the final lamellae were again imaged by iFLM to identify regions for cryo-ET acquisition. The grids with the lamellae were then transferred to a Krios G4 C-FEG cryo-TEM operated at 300 kV for cryo-EM and cryo-ET imaging. Z-stacks acquired by iFLM were correlated to cryo-EM overview maps to target regions containing HRS or EGF signals by cryo-ET. We thereby obtained 5 tomograms with EGF-Alexa647, and 6 tomograms with mScarlet-I3-HRS signals. In one additional tomogram, an EGF-Alexa647 signal was too weak to unambiguously identify, yet the coat was visible. In two of the mScarlet-I3-HRS tomograms, the coat was not identifiable, possibly due to the oblique orientation of membranes relative to the imaging plane. The final data set thus consisted of 10 tomograms, some containing multiple endosomes with visible coat (Figures 4A-C and S7E-P).

Cryo-EM and ET data was acquired on a Krios G4 C-FEG equipped with a Falcon 4 detector used in counting mode and a Selectris energy filter. Data was collected with Tomography 5 Software (Thermo Fisher Scientific). Lamellae maps were acquired at a defocus of −60 µm and a pixel size of 53.7 Å. Tilt series were acquired with a dose-symmetric tilt scheme (Hagen et al., 2017) in groups of 4 between −60° and 60° at an increment of 1° and a pixel size of 2.97 Å. Some tilt series were not acquired at the full range between −60° and 60°. A dose of approximately 1 e-/Å^2^ was applied per tilt. The target defocus was −5 μm. Tilt series alignment was performed with IMOD using patch tracking, and for figure presentation, tomograms were reconstructed by SIRT at bin 2 using IMOD (Mastronarde and Held, 2017; Kremer et al., 1996). A 3D median filter was applied to the reconstructed tomograms for figure presentation.

### Subtomogram averaging

For subtomogram averaging, tomograms were reconstructed by weighted back projection after CTF-correcting tilt series using phase flipping in IMOD (Mastronarde and Held, 2017; Kremer et al., 1996). Of the 12 tomograms with the visible coat, we used 8 for further processing (Table S3).

Subtomogram averaging was performed using Dynamo (Castaño-Díez et al., 2012). Initially, coat-containing membranes were manually annotated using the “surface model” of Dynamo Catalogue (Castaño-Díez et al., 2017) and 1809 segments were extracted on the hard drive. For the initial alignment, a subset of 50 particles was manually coarsely aligned using **dynamo_gallery**. These particles underwent an initial alignment process, and the resulting average was used as a reference for the global alignment of all subtomograms. Subsequent refinement steps involved iterative rounds of alignment and classification in Dynamo. Multireference alignment was performed to sort particles into the classes with the layers and the suboptimal classes. Particles from both “good” and “bad” classes were manually inspected in **dynamo_gallery** to recover misclassified particles. After several rounds of classification and manual inspection, 440 particles were identified as the final set and used for the subtomogram averaging. The final 3D map was first refined without symmetry constraints. Next, disc-like masks were tested for subtomogram averaging, allowing the particles to rotate only around the vertical axis and to shift in-plane. The alignment focused on the first layer resulted in an apparent hexametric lattice. To further enhance the structural details, C6 symmetry was applied in the final rounds of refinement, resulting in a final resolution of 44 Å (Table S3). Similar focused refinements on the other layers did not reveal apparent symmetry.

### Immunofluorescence staining and imaging

For immunofluorescence microscopy, we cultured cells on Poly-D-Lysine (# A3890401, Thermo Fisher Scientific) coated coverslips, washed cells two times with PBS and then fixed them for 15 minutes in room temperature with 4% formaldehyde solution in PBS. Cells were washed three times with 0.2% saponin in PBS, then permeabilized and blocked with 3% BSA, 100 mM glycine, and 0.2% saponin in PBS for 1 hour and then incubated overnight with primary antibodies (see Table S4 for dilutions) in 1% BSA, 0.2% saponin in PBS at +4°C. Cells were then washed three times with saponin-PBS solution and incubated for 1 hour at room temperature with secondary antibodies (Table S4) in 1% BSA 0.2% saponin in PBS. Before mounting the coverslips on microscope slides with ProLong Glass Antifade Mountant (# P36982, Thermo Fisher Scientific), cells were washed two times with PBS and once with water. We imaged cells with a Stellaris 8 Falcon (Leica) confocal microscope equipped with Lightning using a 60x oil objective.

Filipin staining for HeLa mScarlet-I3-HRS cells cultured on Poly-D-lysine coated coverslips was done after cells were fixed for 15 minutes with 4% formaldehyde in PBS. Cells were then washed three times with PBS, and 50 µg/ml Filipin (# F9765, Sigma-Aldrich) in PBS was incubated on cells for 30 minutes at room temperature. Samples were mounted on microscope slides with ProLong Glass Antifade Mountant, and samples were imaged during the same day with a Stellaris 8 Falcon (Leica) confocal microscope equipped with Lightning using a 60x oil objective.

### High-content imaging

To modify cholesterol amounts in the endo-lysosomal pathway, cells were treated with U18666A (# 3633, Sigma-Aldrich) or Methyl-b-cyclodextrin (MbCD, # C4555, Sigma-Aldrich). For the U18666A treatment, cells were cultured on normal media. The following day, 3 µg/ml of U18666A in normal cell culture media was added to cells, and cells were cultured for 24 hours before they were fixed with 4% formaldehyde in PBS. Cholesterol amounts at the endo-lysosomal pathway were verified with Filipin staining.

For high-content imaging, cells were cultured in µ-Plate 96 well plates (# 89626, Ibidi GmbH) and stained as described above. Hoechts 33342 staining (1:10,000 dilution, # H3570, Thermo Fisher Scientific) was performed at the same time with a secondary antibody staining. Cells were imaged with ImageXpress automated spinning disk confocal microscope (Molecular Devices) equipped with an sCMOS camera. Cells were segmented and analyzed with ImageXpress (Molecular Devices). Final graphs were generated with OriginPro 2024b v10.1.5.132.

### Epidermal growth factor (EGF) pulse-chase

We performed the EGF pulse-chase experiment as described in (Wenzel et al., 2018). Shortly, HeLa mScarlet-I3-HRS cells were cultured on Poly-D-lysine coated coverslips, and 50 ng/ml AlexaFluor-647 EGF (# E35351, Thermo Fisher Scientific) in normal cell culture media was incubated with cells for 2 minutes in +37°C. Cells were then washed twice with normal cell culture media and incubated for 5, 15, 30, and 45 minutes in normal cell culture media at 37°C before the fixation with 4% formaldehyde solution in PBS. Cells were then washed three times with PBS before coverslips were mounted on microscope slides as described above. Confocal imaging was done as described above, and colocalization of AlexaFluor-646 EGF with mScarlet-I3-HRS was analyzed with Fiji v1.54f using BIOP JaCoP plugin (https://github.com/BIOP/ijp-jacop-b). Final graphs were generated with OriginPro 2024b v10.1.5.132.

### Cholesterol clustering in large unilamellar vesicles (LUVs)

LUVs were prepared by mixing lipids at 1 mg/ml final concentration in chloroform with a composition of 29 mol% DOPC, 20 mol% DOPE, 20 mol% DOPS, 1 mol% PI(3)P, 29.5 mol% cholesterol, and 0.5 mol% TopFluor-cholesterol. Chloroform was evaporated under argon gas and then in a vacuum, and lipids were hydrated in a reaction buffer (20 mM HEPES pH 7.2, 125 mM KAc). Liposomes were then freeze-thawed 10 times with a water bath and liquid nitrogen and then extruded through 100-nm polycarbonate filters using a mini extruder (Avanti Polar Lipids). To measure TopFluor-cholesterol fluorescence quenching, the final concentration of 50 µM liposomes was mixed with the desired concentration of HRS in the reaction buffer. After 60 minutes, we measured TopFluor-cholesterol spectra between 490 nm and 550 nm with 2 nm steps using a BioTek Synergy H1 plate reader (Agilent Technologies).

### Cholesterol sorting and diffusion with SLBs

SLBs with lipid composition of 29 mol% DOPC, 20 mol% DOPE, 20 mol% DOPS, 1 mol& PI(3)P, 29 mol% cholesterol and 1 mol% TopFluor-cholesterol were prepared as above. HRS at µM concentration and clathrin at 200 nM concentration were incubated with SLBs as described above. We recorded TopFluor cholesterol sorting with a Nikon Eclipse C1 spinning disk confocal microscope using a 100x 1.3 NA objective. The microscope was controlled with Slidebook 6.0.22 (3i Intelligent Imaging Innovations), and the data analysis was performed using Fiji v1.54f. Final graphs were generated with OriginPro 2024b v10.1.5.132.

To measure TopFluor-cholesterol diffusion with fluorescence recovery after the photobleaching (FRAP) approach, we prepared samples with either 500 nM or 2 µM HRS and 200 nM clathrin, as described above. We performed FRAP experiments with an Olympus IX83 TIRF microscope equipped with ImageEM X2 EM-CCD camera (Hamamatsu) and FRAP laser, using Olympus Uapo N 100x NA 1.49 objective. The microscope was controlled with Visiview v.4.4.0.11 software (Visitron Systems). We imaged a timelapse series of SLBs with a 500 ns frame rate. Before bleaching the small region of SLB, we recorded 3-5 frames, and after bleaching, we recorded 1-2 minutes until the recovery plateaued. We analyzed the data with Fiji v1.54f. Final graphs were generated with OriginPro 2024b v10.1.5.132. FRAP immobile fractions and halftimes of mobile fractions were calculated with a one-phase exponential equation:

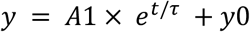

where A is the value of y in the plateau, t is time, τ is a time constant, and y0 is the value of y when t=0. Recovery halftimes were then calculated with the equation:

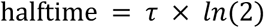

### Cargo protein reconstitution and clustering

LUVs with and without cholesterol (29.9 mol% DOPC, 20 mol% DOPE, 20 mol% DOPS, 1 mol& PI(3)P, 30 mol% cholesterol and 0.1 mol% Atto647-DOPE and 69.9 mol% DOPC, 20 mol% DOPE, 20 mol% DOPS, 1 mol& PI(3)P, and 0.1 mol% Atto647-DOPE), respectively, were used to reconstitute SfGFP-VAMP2-4xUb produced recombinantly, as previously performed with other transmembrane proteins (Espadas et al., 2019)(Figure 6A). Briefly, LUVs were initially incubated in the presence of 0.1% Triton ×100 for 30 minutes (Figure 6A, step 1). Detergent-destabilized liposomes were then mixed with purified protein at a protein:lipid molar ratio of 1:500. After 30 minutes of incubation at room temperature with gentle agitation, the Triton ×100 detergent was removed using BioBeads SM-2 adsorbent (BioRad) by four consecutive additions to the proteo-lipid mixture every hour, followed by an overnight incubation (Rigaud et al., 1995). The sample was then spin-down to remove the BioBeads and non-incorporated protein, and the supernatant containing the proteoliposomes was collected (Figure 6A, step 2).

To perform the GUVs with reconstituted protein, proteoliposomes were dialyzed against a 25 mM HEPES and 1 mM Trehalose solution for 2 hours. After dialysis, 15 µl of the freshly dialyzed sample was placed on a clean parafilm surface and mixed with 2 µl of 40 um silica beads (Microspheres-Nanospheres, USA), and then dried in a vacuum chamber for at least 1 hour (Figure 6A, step 3). The dried beads covered with the proteo-lipid layers were then transferred to a tube containing 10 µl of 1M Trehalose solution for 15 min at 65 °C. Next, the 10 µl containing the pre-hydrated proteo-lipid lamellas were transferred to the observation chamber containing the working buffer (20 mM HEPES, pH 7.2, 125 mM KAc, 1 mM MgAc) for complete proteo-GUVs hydration (Figure 6A, step 4). Lastly, mScarlet-I3-HRS at a final concentration of 400 nM was added to the microscopy chamber and incubated for 30 min before starting the imaging, using a spinning-disk Nikon Ti-Eclipse.

## Author contributions

M.H., S.B.M., I.G., M.G., C.B.S., J.E., C.M., A.C., and S.V. carried out experiments and data analysis. Mi.K., W.K., A.C., S.V., Ma.K., and A.R. contributed to experiment planning and supervised the work. M.H., Ma.K., and A.R. conceived the study and planned experiments. M.H. and A.R. wrote the manuscript with input from all authors.

## Acknowledgments

We are grateful to Jean Gruenberg, Stefania Vossio, and Elina Ikonen for thoughtful discussions about the role of cholesterol in vesicle trafficking. Andrea Picco is acknowledged for technical support in TIRF imaging, and Rafael Ferreira Caetano and Frédéric Humbert for protein purification. We thank Roux and Kaksonen labs for technical support and helpful discussions. We thank the Photonic Light Microscopy Facility and ACCESS Facility at the University of Geneva, the Microscopy Imaging Center (MIC) of the University of Bern, the Dubochet Center for Imaging (DCI) Bern, and IBPS electron microscopy platform at Sorbonne University in Paris for microscope access and support with data collection. This work was supported by EMBO (ALTF 703-2020 for M.H., and 989-2022 for J.E.), Fundación Alfonso Martín Escudero (Postdoctoral fellowship for C.B-S.), and Swiss National Science Foundation grant (310030_212288) for Ma.K. A.R. acknowledges funding from the Swiss National Fund for Research grant numbers #CRSII5_189996 and #310030_200793 and the European Research Council Synergy grant number #951324-R2-TENSION. Work in the group of W.K. was supported by the University of Bern and the SNSF project 201158. We would also like to acknowledge funding by the *Agence Nationale pour la Recherche* (ANR-20-CE13-0024-01, ANR-21-CE13-0018-01 to SV), Horizon Europe “DREAMS” project under grant agreement N°101080229, Sorbonne Université, INSERM, Association Institut de Myologie core funding to S.V. Mi.K. is supported by the Helmholtz Society and the Heisenberg Award from the DFG (KU3222/3-1).

