## Supplementary information for "Two-dimensional HRS condensates drive the assembly of flat clathrin lattices on endosomes"

- Figures S1-11
- Tables S1-4
- Supplementary movie legends

### Supplementary figures

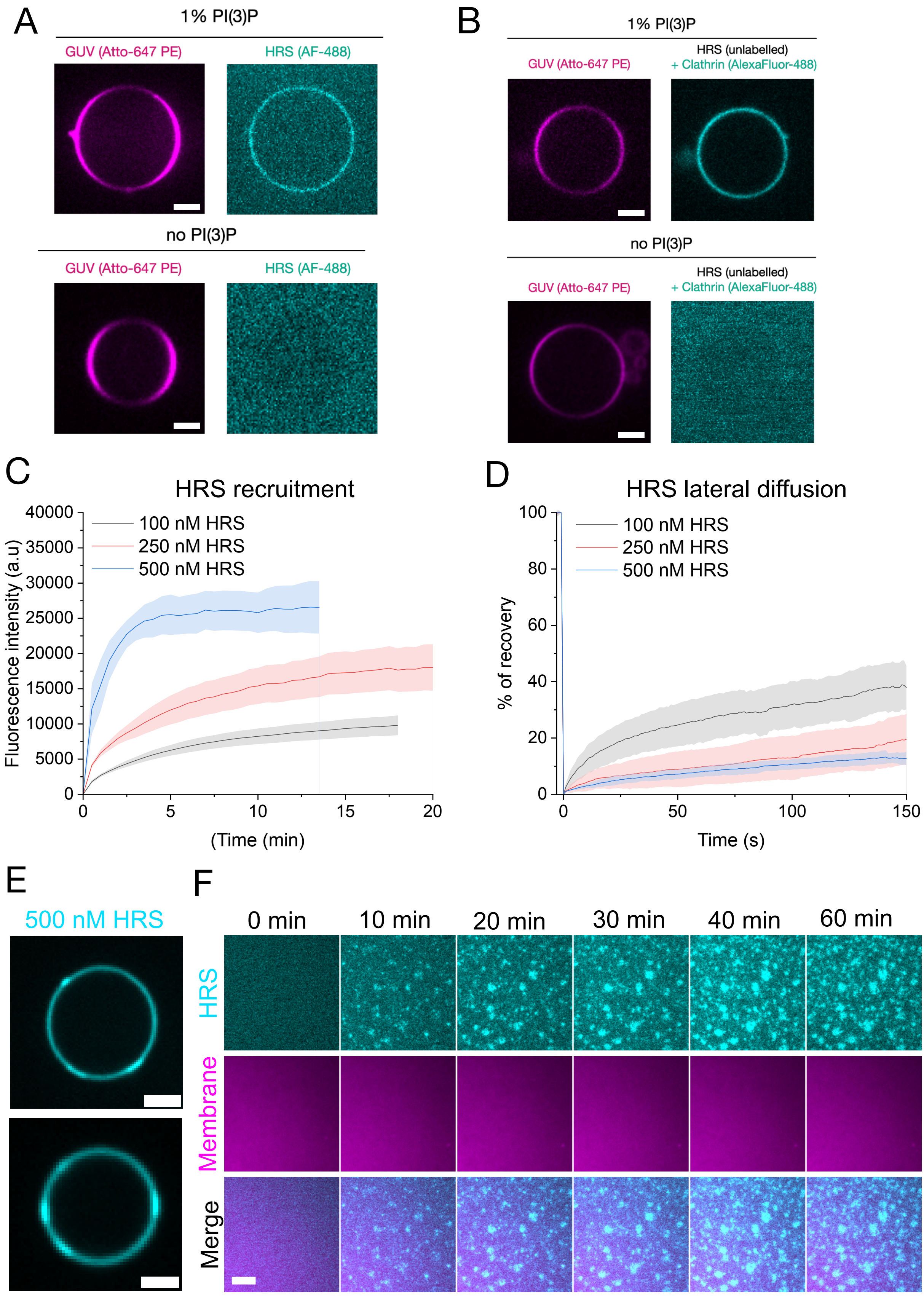

**Figure S1**

(A) Representative images of 250 nM AlexaFluor-488 labelled HRS binding on GUVs with either 1% PI(3)P or without PI(3)P. (B) Representative images of AlexaFluor-488 labeled clathrin recruitment on PI(3)P-rich or PI(3)P-deficient GUVs in the presence of unlabeled HRS. Scale bars are 5 μm. (C) The recruitment of 100 nM, 250 nM, and 500 nM HRS on 1% PI(3)P-rich SLB. Data is a mean of four measurements with a standard deviation shown. The experiment was repeated two times with similar results. (D) FRAP of AlexaFluor-568 labeled HRS on the SLB. Data is a mean of 20 measurements (100 nM HRS), 18 measurements (250 nM HRS), and eight measurements (500 nM HRS) with standard deviations shown. (E) Representative images of GUVs incubated with 500 nM HRS (AlexaFluor-488 labelled). Scale bars are 5 μm. (F) Representative images of SLBs incubated with 500 nM HRS (AlexaFluor-488 labelled). Scale bars are 5 μm.

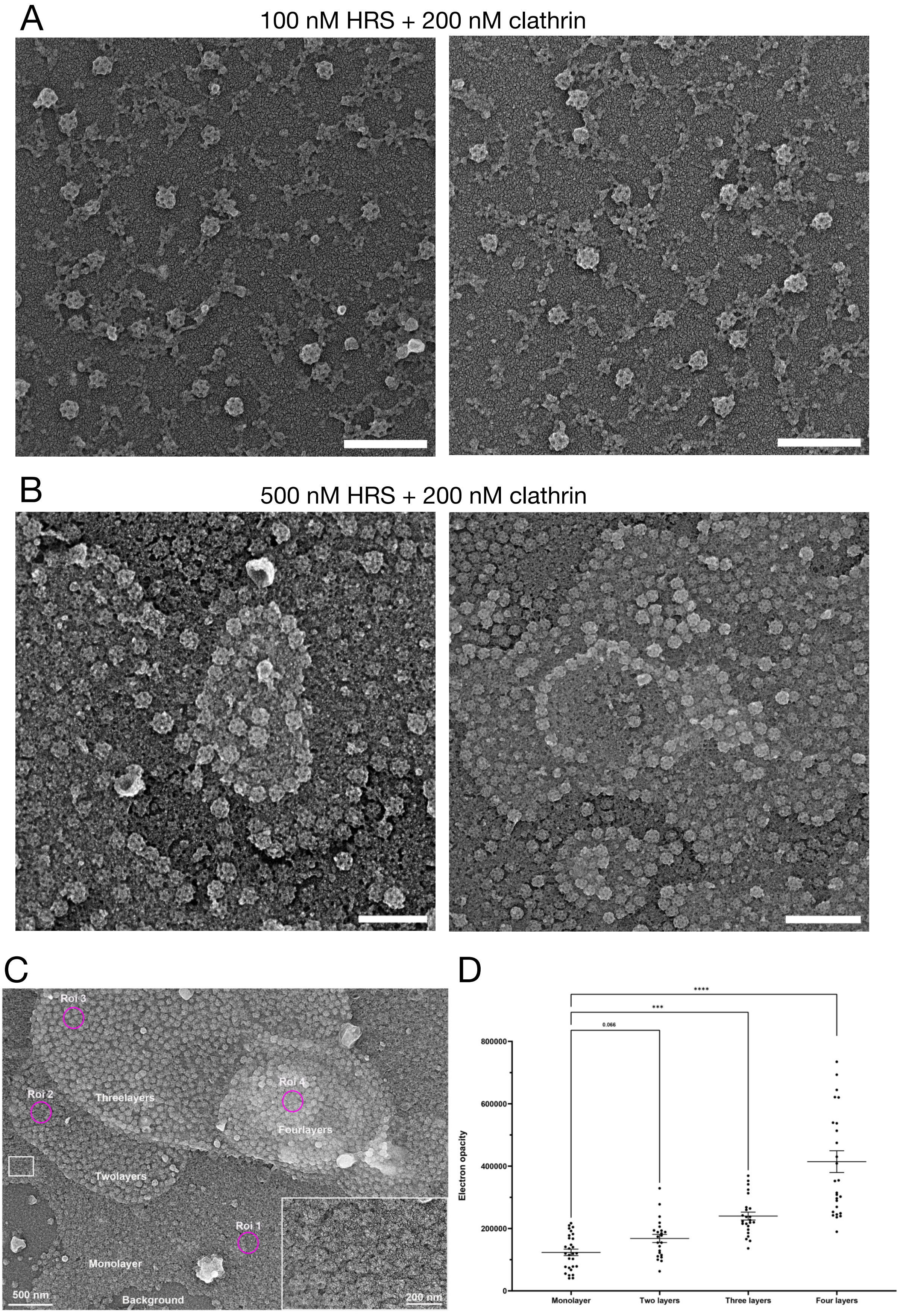
A
100 nM HRS + 200 nM clathrin
B
500 nM HRS + 200 nM clathrin
C
Roi 3
Roi 2
Threelayers
Roi 4
Fourlayers
Twolayers
Roi 1
Monolayer
Background
500 nm
200 nm
D
****
***
0.066
800000
600000
400000
200000
0
Electron opacity
Monolayer
Two layers
Three layers
Four layers

**Figure S2**

Additional examples of platinum replica microscopy images of reconstituted samples with (A) 100 nM HRS and 200 nM clathrin, and (B) 500 nM HRS and 200 nM clathrin. The scale bars are 300 nm. (C) Representative images of reconstituted 1 μM HRS and 200 nM clathrin lattices on supported lipid bilayers (SLBs), captured using PREM. The PREM image reveals clathrin lattices with varying layer numbers (two, three, and four layers). Multilayered structures predominantly show clathrin pit morphology, whereas the monolayer mainly forms a flat clathrin lattice (inset). Scale bars represent 500 nm and 200 nm (inset). (D) Quantification of clathrin layer density on SLBs. Regions of interest (ROIs), indicated by pink circles as shown in panel a, were randomly selected from different layers, and the corresponding electron opacity values were measured. The histogram represents pooled data from multiple ROIs across three independent experiments. Statistical significance was assessed using a two-tailed t-test.

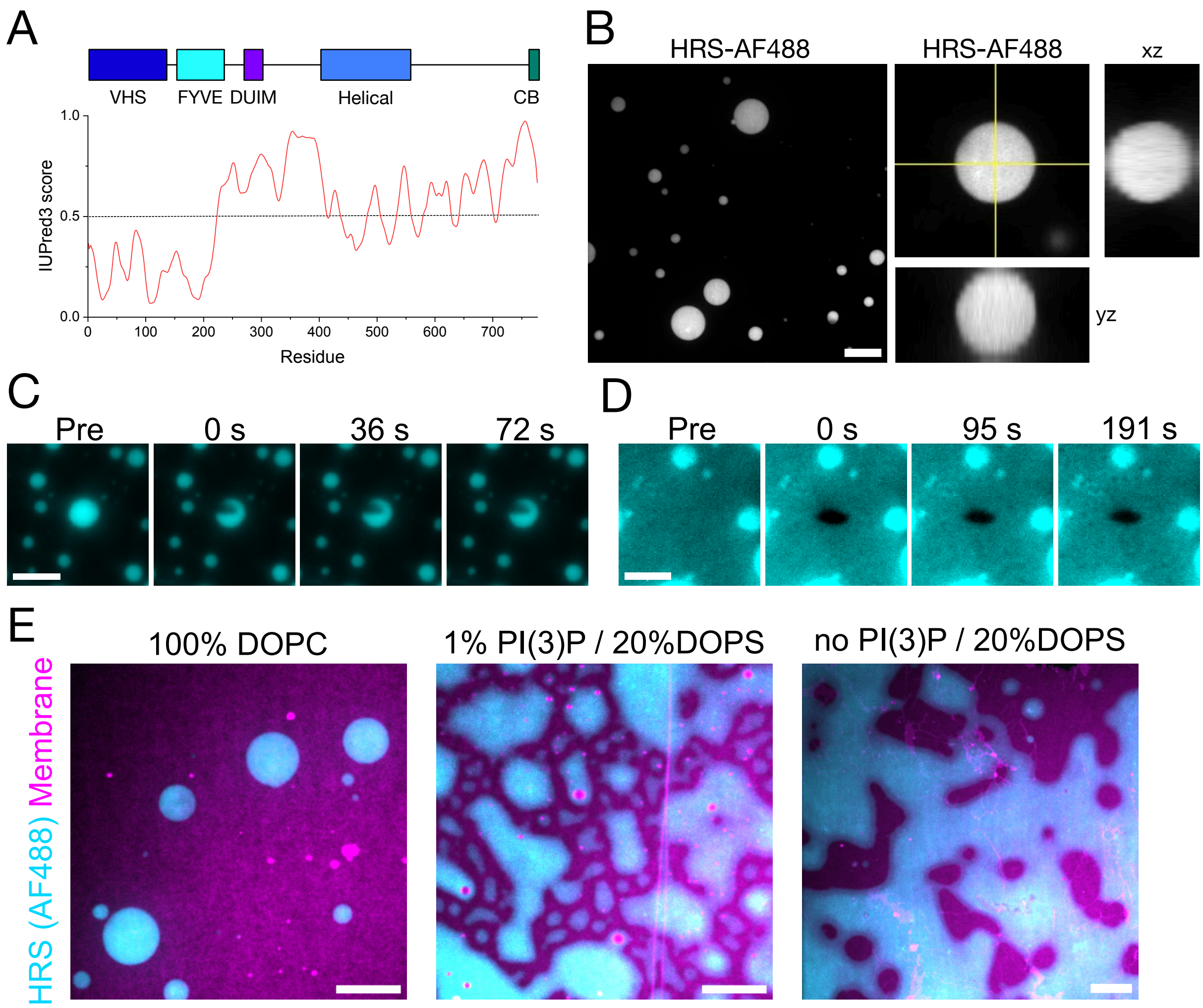


**Figure S3**

(A) Domains and IUPred3 prediction of unstructured regions in HRS. (B) Representative fluorescence microscopy images of 2 µM AlexaFluor-488 labeled HRS in 20 mM HEPES, pH 7.2, 125 mM potassium acetate, and 1 mM magnesium acetate. The scale bar is 10 µm. Orthogonal views of HRS condensate are shown. (C) Representative time-lapse images of FRAP experiments of 2 µM HRS (AlexaFluor-488 labeled) droplet on SLB membrane. The scale bar is 5 µm (D) Representative time-lapse images of FRAP experiments of 2 µM HRS (AlexaFluor-488 labeled) bidimensional condensates on SLB membrane. The scale bar is 5 µm. (E) 2 µM HRS (AlexaFluor-488 labeled) on SLB containing 100% DOPC (left), 29% DOPC, 20% DOPE, 20% DOPS, 1% PI(3)P, and 30% cholesterol (middle), and 30% DOPC, 20% DOPE, 20% DOPS, and 30% cholesterol (right). All membranes are supplemented with 0.05% DOPE Atto-647n. The scale bar is 10 µm.

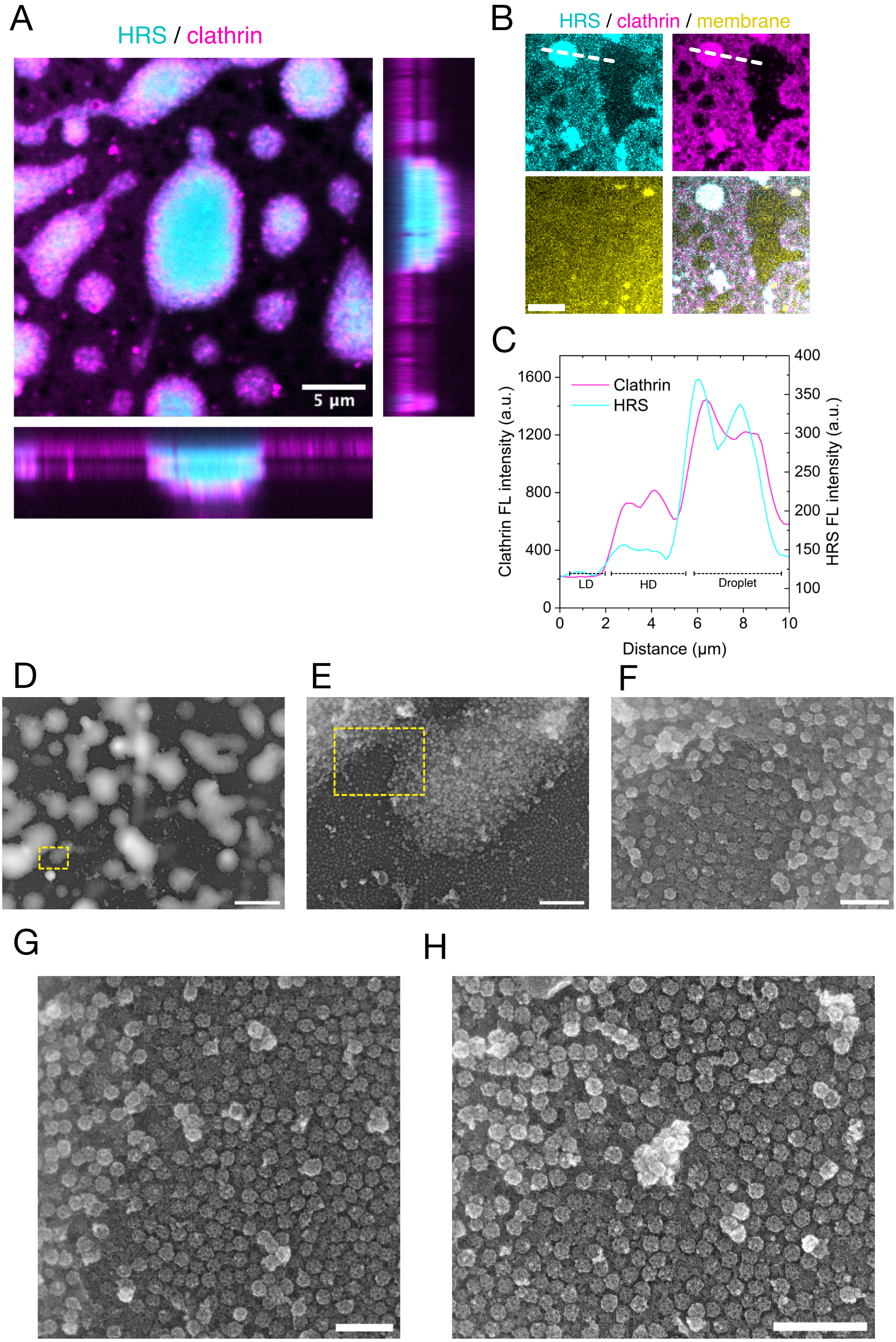
A
HRS / clathrin
5 μm
B
HRS / clathrin / membrane
C
Clathrin
HRS
Clathrin FL intensity (a.u.)
HRS FL intensity (a.u.)
Distance (μm)
LD
HD
Droplet
D
E
F
G
H

**Figure S4**

(A) A representative fluorescence microscopy image of 2 μM HRS (AlexaFluor-488 labeled, cyan) and 200 nM clathrin (AlexaFluor-568 labeled, magenta) on SLB. The scale bar is 5 μm. (B) Representative fluorescence microscopy images of 2 μM HRS (AlexaFluor-488 labeled, cyan) and 200 nM clathrin (AlexaFluor-568 labeled, magenta) on SLB (yellow), forming clathrin-coated two-dimensional condensates. LD=low-density HRS, HD=high-density HRS, D=droplet condensate. The scale bar is 10 μm. (C) Line profile analysis of AlexaFluor-488 labeled HRS and AlexaFluor-568 labeled clathrin over three HRS populations on supported bilayers in panel (B). (D-F) PREM images of clathrin-coated spherical HRS condensates reconstituted with 2 μM HRS and 200 nM clathrin. (G-H) PREM images of clathrin-coated two-dimensional HRS condensates reconstituted with 2 μM HRS and 200 nM clathrin. Scale bars are 10 μm (A), 1 μm (B) and 300 nm (C-E).

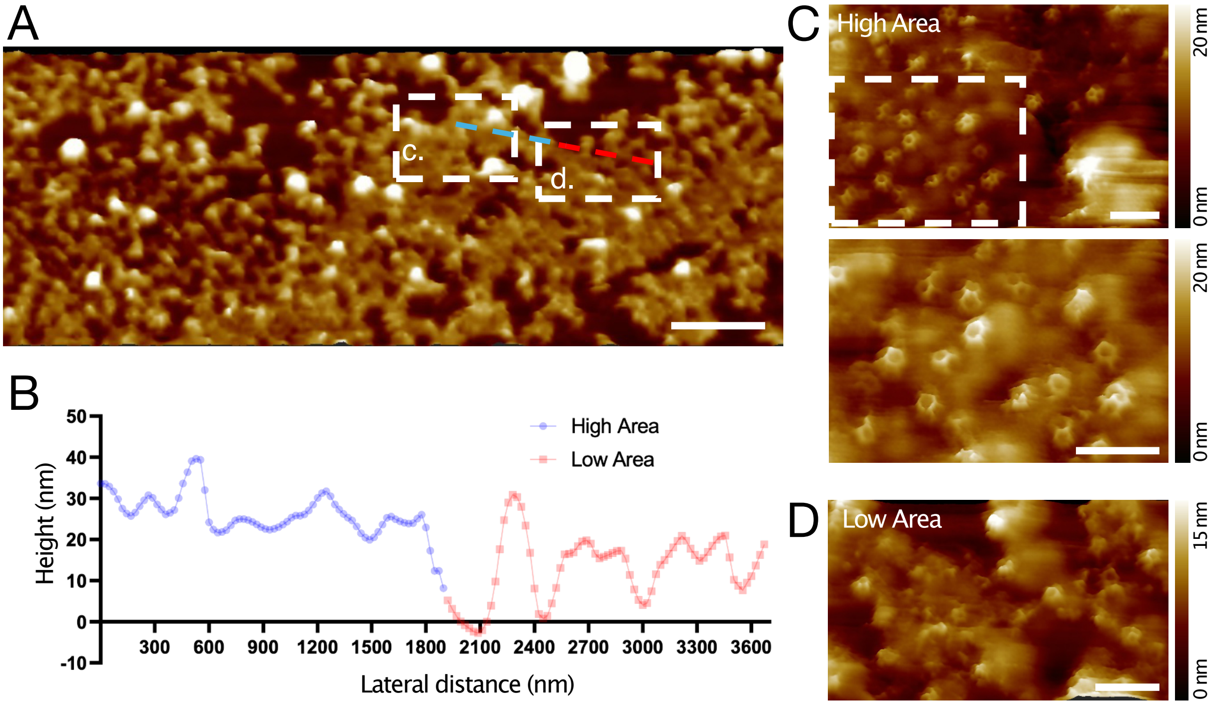


**Figure S5**

(A) A representative HS-AFM image of HRS-clathrin coat on SLB. The scale bar is 2500 nm. Scattered boxes indicate regions where a higher resolution image was acquired for panels (C) and (D). The light blue-red scattered line indicates the region where the line profile was measured for panel (B). (B) A line profile analysis of the height of the HRS-clathrin coat. Blue and red lines represent high region and low coat regions, respectively. (C) A high-resolution image of the clathrin coat at the high coat region. The scattered box indicates the region where the zoom-in image. Scale bars are 200 nm. (D) A high-resolution image of the clathrin coat on the low coat region. A scale bar is 200 nm.

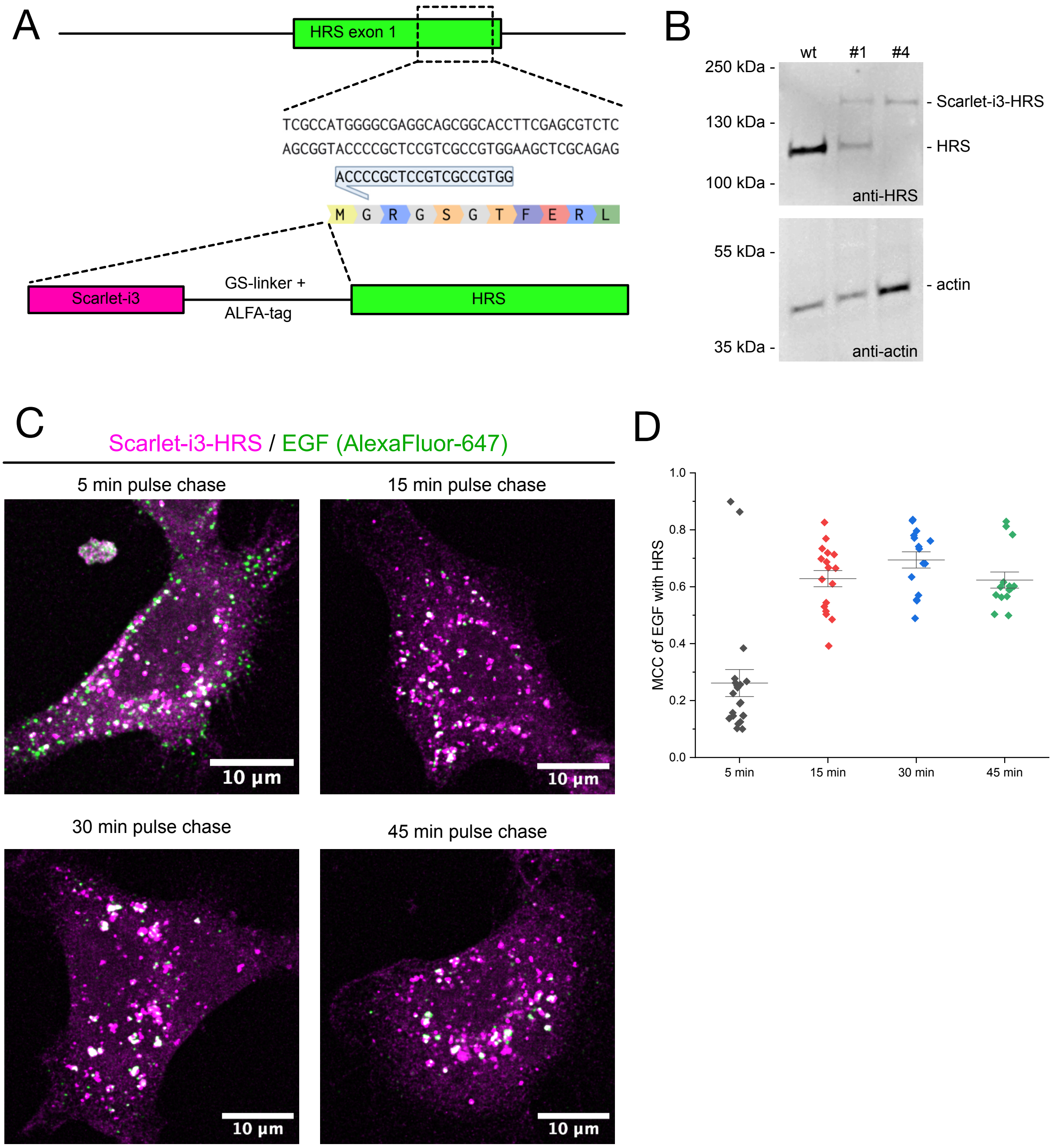


**Figure S6**

(A) A schematic presentation of CRISPR-Cas9 knock-in strategy (see Methods for details). (B) Western blot of control HeLa cells and CRISPR-Cas9 mScarlet-I3-HRS knock-in cell lines. (C) Representative fluorescence microscopy images of mScarlet-I3-HRS (magenta) cell lines with 5 min, 15 min, 30 min, or 45 min pulse of EGF-AlexaFluor-647 (green). Scale bars are 10 µm. (D) Mander's colocalization coefficients of EGF-AlexaFluor-647 colocalizing with HRS.

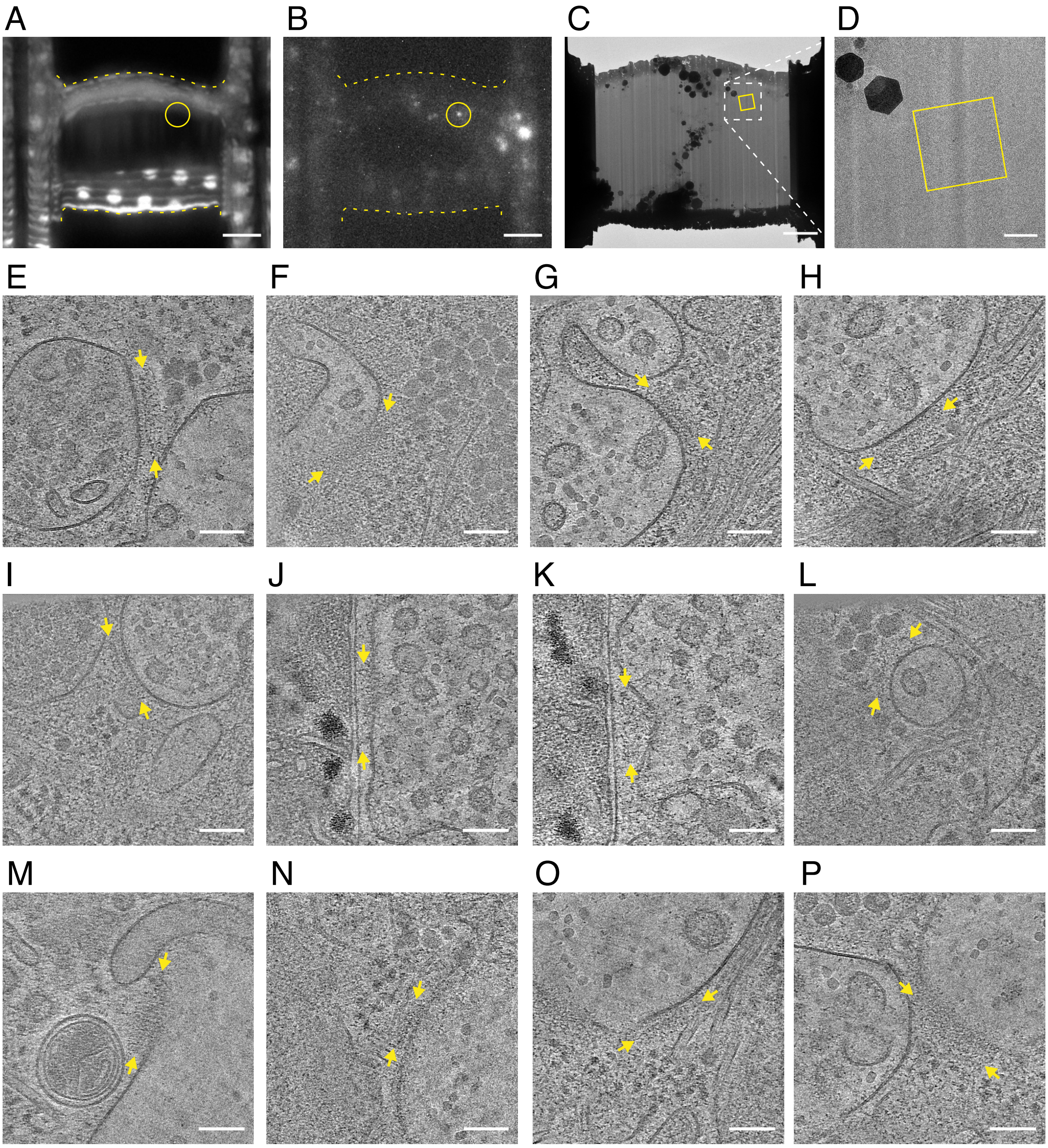


**Figure S7**

(A-D) The cryo-CLEM workflow for the acquisition of the data shown in Figure 4A-C. (A) An image of the lamella after rough milling, acquired by iFLM in reflection mode. The outline of the lamella is indicated by the yellow dashed line. (B) A fluorescence image of the same lamella acquired by iFLM in fluorescence mode, showing the signal of AlexaFluor-647 EGF. The yellow circle indicates the target signal. (C) A cryo-EM overview of the final lamella. The yellow square indicates the area of acquisition of the tomogram shown in Figure 4A. (D) Zoom into cryo-EM overview, as indicated by the white dashed square in panel (C). (E-P) All regions with a putative protein coat on endosomes (yellow arrows) identified in tomograms acquired in a similar manner as indicated by the workflow in panels (A-D) and Figure 4A. Scale bars are 5 µm (A-B), 3 µm (C), 500 nm (D), and 100 nm (E-P).

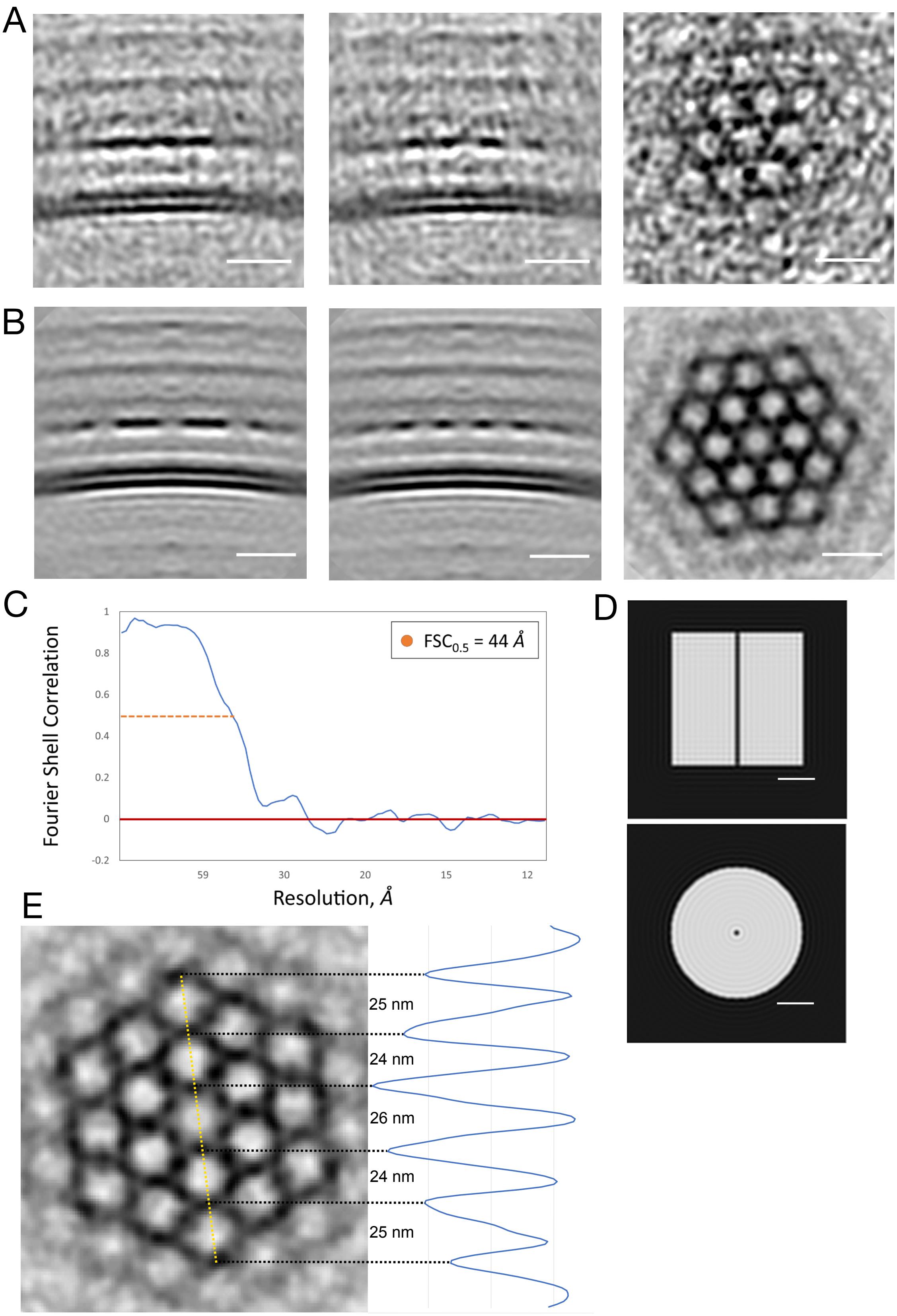

A
B
C
Fourier Shell Correlation
FSC$_{0.5}$ = 44 Å
1
0.8
0.6
0.4
0.2
0
-0.2
59
30
20
15
12
Resolution, Å
D
E
25 nm
24 nm
26 nm
24 nm
25 nm

**Figure S8**

(A) Side view (XZ and YZ) and top view of the initial structure. Scale bars are 20 nm for side-view images and 40 nm for top-view images. (B) Side view (XZ and YZ) and top view of the refined structure. Scale bars are 20 nm for side-view images and 40 nm for top-view images. (C) A Fourier Shell Correlation (FSC) graph. (D) The mask used for the refinement. (E) The analysis of vertex-to-vertex distance of hexagonal lattice.

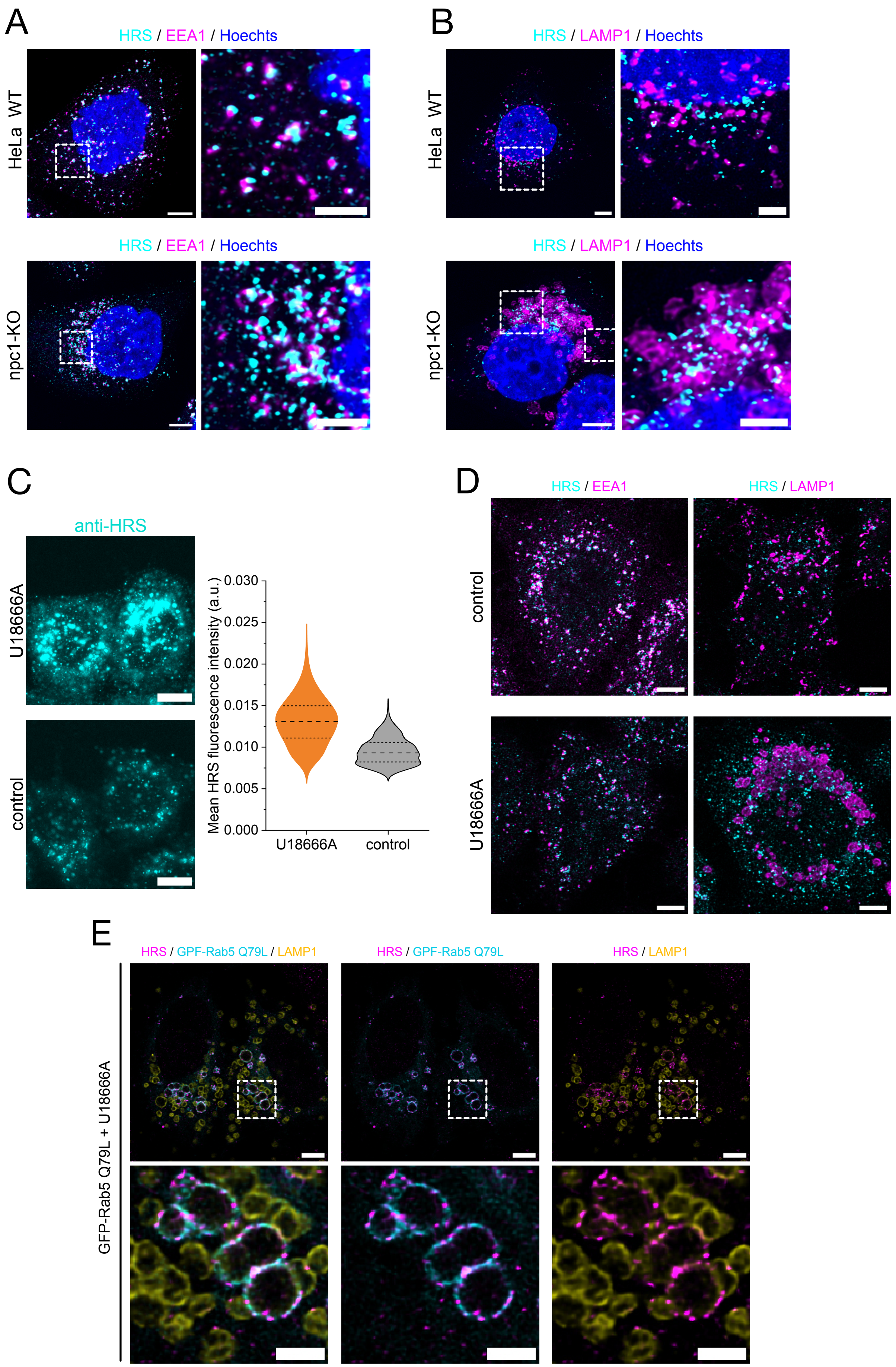
A
HRS / EEA1 / Hoechts
HeLa WT
HRS / EEA1 / Hoechts
npc1-KO
B
HRS / LAMP1 / Hoechts
HeLa WT
HRS / LAMP1 / Hoechts
npc1-KO
C
anti-HRS
U18666A
control
Mean HRS fluorescence intensity (a.u.)
0.030
0.025
0.020
0.015
0.010
0.005
0.000
U18666A
control
D
HRS / EEA1
HRS / LAMP1
control
U18666A
E
HRS / GPF-Rab5 Q79L / LAMP1
HRS / GPF-Rab5 Q79L
HRS / LAMP1
GFP-Rab5 Q79L + U18666A

**Figure S9**

(A) A representative fluorescence microscopy image of HeLa wild type and NPC1 knockout cells stained with HRS (cyan) and EEA1 (magenta) antibodies. (B) A representative fluorescence microscopy image of HeLa wild type and NPC1 knockout cells stained with HRS (cyan) and LAMP1 (magenta) antibodies. In panels A-B, scale bars are 5 μm for larger images and 2 μm for zoom-in images. (C) Representative fluorescence microscopy images of HeLa control cells and cells treated with U18666A stained with HRS antibody (cyan). Scale bars are 10 μm. Mean HRS fluorescence intensity in U18666A treated and control HeLa cells. Experiments was repeated three times (N=3). (D) Representative images of control HeLa cells and U18666A drug-treated cells with HRS, EEA1, and LAMP1 immunofluorescence staining. Scale bars are 20 μm. (E) A representative fluorescence microscopy image of HeLa cells transfected with GFP-Rab5 Q79L mutant (cyan) and treated with U18666A, and stained HRS (magenta) and LAMP1 (yellow) antibodies. Scale bars are 5 μm for larger images and 2 μm for zoom-in images.

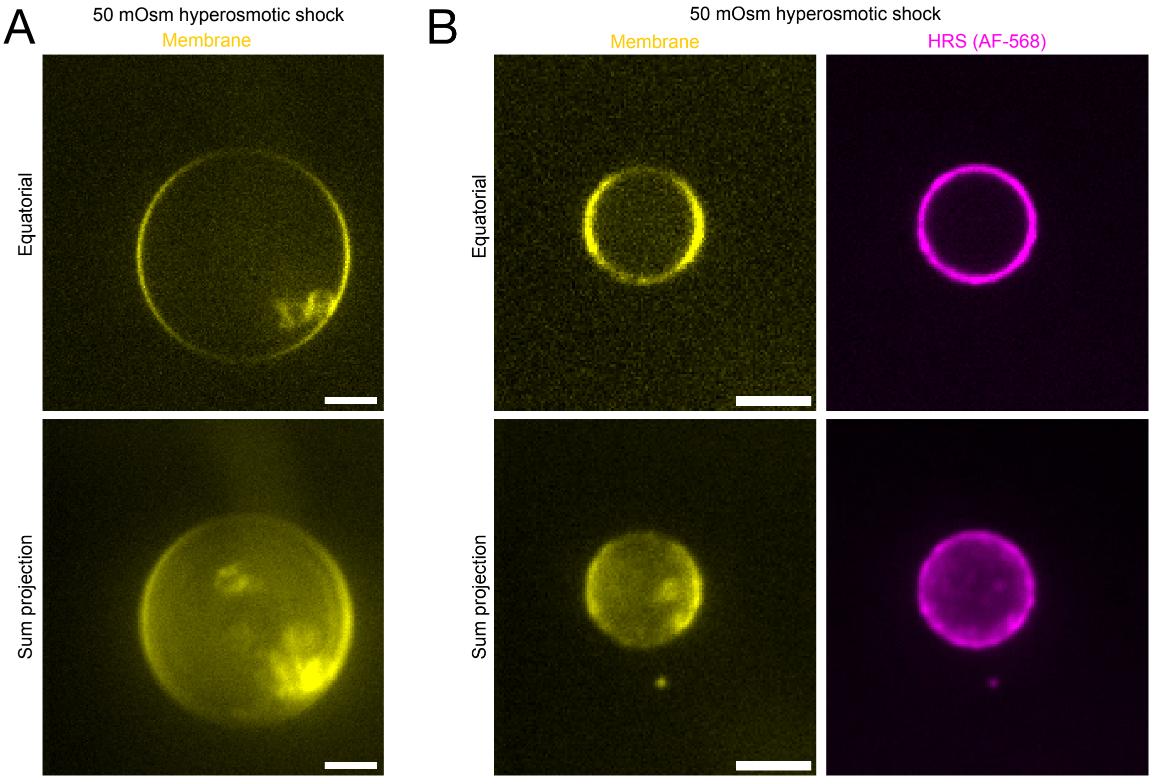


**Figure S10**

(A) GUV (yellow) containing 30% cholesterol under hyperosmotic shock. The equatorial plane and sum projection are shown. (B) GUV (yellow) containing 30% cholesterol incubated with 500 nM HRS (AlexaFluor-568 labeled, magenta) under hyperosmotic shock. The equatorial plane and sum projection are shown. Scale bars are 5 μm.

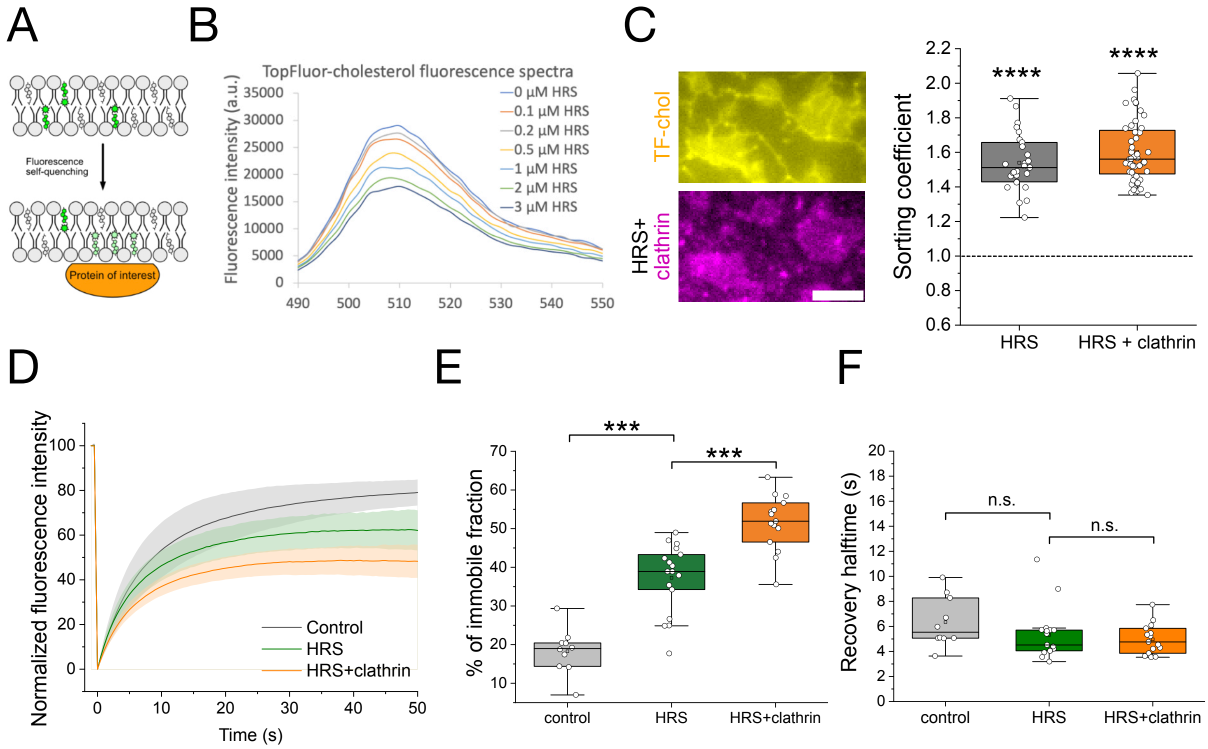


**Figure S11**

(A) A schematic presentation of fluorescence quenching assay. (B) TopFluor-cholesterol fluorescence spectra with different HRS concentrations. The experiment was repeated three times (N=3) with similar results. (C) TopFluor-cholesterol sorting coefficient analysis with HRS or HRS and clathrin. A representative fluorescence microscopy image of 1% TopFluor-cholesterol (yellow) in SLB incubated with 2 µM HRS (unlabeled) and 200 nM clathrin (20% AF-568 labeled, magenta). The scale bar is 5 µm. Sorting coefficient values were blotted for 2 µM HRS alone and 2 µM HRS + 200 nM clathrin samples. Each data point represents sorting coefficient measurements on independent SLB from three experiments. (D-F) FRAP experiments to measure TopFluor-cholesterol diffusion in control samples (no proteins), with 500 nM HRS, and with 500 nM HRS and 200 nM clathrin. (D) Recovery curves of TopFluor-cholesterol after photobleaching in control samples (grey) and in the presence of HRS (green), and HRS and clathrin (orange). Dark lines represent mean values and shaded areas standard deviations. (E) Percentage of immobile fraction of TopFluor-cholesterol FRAP experiment and (F) recovery halftimes of TopFluor-cholesterol FRAP experiments in control samples (no proteins), with 500 nM HRS, and with 500 nM HRS and 200 nM clathrin. In panels (E-F), each data point represents an independent measurement from in total of three experiments (N=3). Statistical testing: Kruskal-Wallis test, n.s. = not statistically significant, *** $p < 0.001$, **** $p < 0.0001$.

#### Supplementary tables

**Table S1.** Lipids used in this study. Their abbreviation, full name, commercial source and catalog numbers are mentioned.

| Abbreviation | Full name | Source | Catalog # |
|---|---|---|---|
| DOPC | 1,2-dioleoyl-sn-glycero-3-phosphocholine | Avanti Polar Lipids | 850375 |
| DOPE | 1,2-dioleoyl-sn-glycero-3-phosphoethanolamine | Avanti Polar Lipids | 850725 |
| DOPS | 1,2-dioleoyl-sn-glycero-3-phospho-L-serine | Avanti Polar Lipids | 840035 |
| PI(3)P | 1,2-dioleoyl-sn-glycero-3-phospho-(1'-myo-inositol-3'-phosphate | Avanti Polar Lipids | 850150 |
| Cholesterol | cholesterol (plant) | Avanti Polar Lipids | 700100 |
| TopFluor-cholesterol | 23-(dipyrrometheneboron difluoride)-24-norcholesterol | Avanti Polar Lipids | 810255 |
| DOPE Atto-647n | Atto 647n 1,2-Dioleoyl-sn-glycero-3-phosphoethanolamine | Attotec | AD 647N-161 |

**Table S2.** Lipid compositions used in this study. Lipid mix number as referred in text, key features, and full composition with molar percentages are listed.

| Lipid mix # | Key features | Full composition | Mol% |
|---|---|---|---|
| # 1 | 1% PI(3)P / 15% cholesterol | DOPC:DOPE:DOPS:PI(3)P:cholesterol:DOPE Atto-647n | 44:20:20:1:15:0.05 |
| # 2 | 0% PI(3)P, 20% DOPS | DOPC:DOPE:DOPS:cholesterol:DOPE Atto-647n | 45:20:20:15:0.05 |
| # 3 | 1% PI(3)P / 0% cholesterol | DOPC:DOPE:DOPS:PI(3)P: DOPE Atto-647n | 56:20:20:1:0.05 |
| # 4 | 1% PI(3)P / 30% cholesterol | DOPC:DOPE:DOPS:PI(3)P:cholesterol:DOPE Atto-647n | 29:20:20:1:30:0.05 |
| # 5 | 1% PI(3)P / 30% cholesterol / 1% TF-cholesterol | DOPC:DOPE:DOPS:PI(3)P:cholesterol:TopFluor-cholesterol:DOPE Atto-647n | 29:20:20:1:29:1:0.05 |
| # 6 | 1% PI(3)P / 30% cholesterol / 0.1% TF-cholesterol | DOPC:DOPE:DOPS:PI(3)P:cholesterol:TopFluor-cholesterol:DOPE Atto-647n | 29:20:20:1:30:0.1:0.05 |
| # 7 | 100% DOPC | DOPC:DOPE Atto-647n | 100:0.05 |

**Table S3** Data collection and processing statistics for the StA

| | |
|---|---|
| Microscope | Titan Krios G4 |
| Magnification | 42,000 |
| Voltage (kV) | 300 |
| Cs (mm) | 2.7 |
| Total electron dose ($e^-$/Å$^2$) | ≤ 120 |
| Defocus range (µm) | 3.5 – 4.1 (determined with ctfplotter) |
| Camera | Falcon 4 with Selectris energy filter |
| Pixel size (Å) | 2.97 (data collection)<br>5.94 (processing) |
| Number of tomograms | 8 |
| Symmetry imposed | C6 |
| Initial number of particles | 1809 |
| Final number of particles | 440 |
| Refinement method | Non-independent half-sets |
| Map resolution (Å) | 44 |
| FSC threshold | 0.5 |

**Table S4.** Antibodies and dilutions used in immunofluorescence imaging (IF) and in Western blot (IB) in this study.

| Target | Antibody name | Source | Catalog number | Host species | Dilution in IF | Dilution in WB |
|---|---|---|---|---|---|---|
| HRS | Anti-HGS | Abcam | Ab72053 | Rabbit | 1/300 | 1/1000 |
| EEA1 | Anti-EEA1 | BD Biosciences | 610457 | Mouse | 1/300 | - |
| LBPA | LBPA antibody serum | Jean Gruenberg lab, University of Geneva | - | Mouse | 1/100 | - |
| LAMP1 | Anti-LAMP1 (H4A3) | Developmental Studies Hybridoma Bank (DSHB) | H4A3 | Mouse | 1/500 | - |
| Actin | Anti-actin | Sigma-Aldrich | A2066 | Rabbit | - | 1/1000 |

#### Supplementary Movie legends

**Movie S1**
A PREM tomography of HRS- clathrin reconstitution on SLB with 500 nM HRS and 200 nM clathrin.

**Movie S2**
An example of fusion events of 2 µM AlexaFluor-488 -labeled HRS condensates in 20 mM HEPES, pH 7.2, 125 mM KAc, 1 mM MgAc buffer.

**Movie S3**
An example of 2 µM AlexaFluor-488 -labeled HRS condensates (cyan) spreading on SLB (not shown)

**Movie S4**
An example of HRS recruitment on SLB membrane acquired with HS-AFM. The scale bar is 1 µm.

**Movie S5**
An example of clathrin recruitment on HRS-rich SLB membrane acquired with HS-AFM. The scale bar is 500 nm.

**Movie S6**
An example of 4xUb-VAMP2-sfGFP (cyan) clustering in GUV membrane containing 0% cholesterol. The sample was incubated 500 nM mScarlet-I3-HRS (magenta)

**Movie S7**

An example of 4xUb-VAMP2-sfGFP (cyan) clustering in GUV membrane containing 30% cholesterol. The sample was incubated 500 nM mScarlet-I3-HRS (magenta)
